# Reproductive output of old polygynous males is limited by seminal fluid, not sperm number

**DOI:** 10.1101/2024.07.02.601670

**Authors:** Krish Sanghvi, Sucheta Shandilya, Alana Brown, Biliana Todorova, Martin Jahn, Samuel J.L. Gascoigne, Tara-Lyn Camilleri, Tommaso Pizzari, Irem Sepil

**Affiliations:** Department of Biology, 11a Mansfield Road, University of Oxford. OX13SZ. U.K.

**Keywords:** Ageing, cryptic female choice, ejaculate, senescence, sexual conflict, sexual selection

## Abstract

Advancing male age can lead to reproductive senescence, which in males is thought to be largely driven by declines in the numbers of sperm transferred by old males. This decline is predicted to be particularly pronounced in polygynous species, where males become sperm limited over a mating sequence. However, males also transfer seminal fluid to females, and little is known about the contribution of seminal fluid to constrain the reproductive output of old, multiply-mating males. Using *Drosophila melanogaster*, we investigated whether age-related variation in male reproductive output is driven by differential limitation of sperm or seminal fluid, over a series of experimental matings. Consistent with reproductive senescence, old males produced fewer offspring than young males. However, this pattern was not driven by sperm limitation, with old males having more sperm and transferring similar numbers of sperm to a female, compared to young males. Yet surprisingly, females stored fewer sperm when mated to old than young males. Notably, females mated to old multiply-mating males produced more offspring when supplemented with seminal fluid, suggesting that fertility of old males over successive matings was limited by seminal fluid availability. Generally, our study indicates that germline maintenance might be prioritised over somatic maintenance as hypothesized by the disposable soma theory of ageing, and that seminal fluid senescence is a key contributor of reproductive decline with age. While other factors such as differential sperm viability and female post-mating responses could have also influenced our results, our study highlights the under-appreciated role of seminal fluid in mediating male reproductive senescence in polygynous species.

**Significance statement:** A key assumption in ageing research is that old males are less fertile than young males, and that this reduced fertility is partly driven by old males producing fewer sperm. However, senescence in male fertility can be caused via other ejaculate-mediated pathways, which we investigate using fruit flies. Contrary to expectations, we reveal that senescence in male fertility is not because of declines in male sperm reserves, but is instead due to age-related changes in seminal fluid and differences in female sperm storage with male age. These declines in the reproductive output of old, multiply-mating males are alleviated by supplementing females with “extra” seminal fluid. Our study demonstrates that male reproductive senescence is reversible, highlighting the underappreciated role of seminal fluid in modulating senescence in polygynous species. These results have potential for improving animal fertility and our understanding of sexual selection.

## Introduction

Advancing age, often (Lemaitre and Gaillard, 2017; Monaghan et al, 2008), but not always (Jones et al, 2014; Sanghvi et al, 2024), leads to irreversible declines in fecundity and fertility (Vrtilek et al, 2023), a pattern known as reproductive senescence (Monaghan and Metcalfe, 2019). In males, reproductive senescence is primarily attributed to age-dependent deterioration in mating success (Amin et al, 2012; Rezaei et al, 2015), reduced sperm quality (Dean et al, 2010; Gasparini et al, 2019; Johnson et al, 2015; Vuarin et al, 2019), sperm quantity (Cornwallis et al, 2014; Sasson et al, 2012), and sperm performance (Aich et al, 2021; Gasparini et al, 2010). However, male reproductive senescence can also occur due to age-dependent deterioration in the non-sperm component of the ejaculate, seminal fluid (Borziak, 2016; Fricke and Koppik, 2019; Koppik and Fricke, 2017; Sepil et al, 2020).

Understanding male reproductive senescence requires an integrated study of both, sperm- and seminal fluid-mediated changes in male reproductive output with age (Koppik and Fricke, 2017; Sepil et al, 2020). In many species, seminal fluid is crucial for ensuring fertilization (Chapman and Davies, 2004), maintaining sperm performance (den Boer et al, 2010; Ramm, 2020), promoting female oviposition (Heifetz et al, 2005; Poiani, 2006; Sirot et al, 2009b), and facilitating long-term sperm storage (Hopkins et al. 2017; Avila et al, 2011). Ejaculates produced by old males often have an altered seminal fluid protein (SFP) composition (Koppik and Fricke, 2017, Sepil et al, 2020), lower quantities of seminal fluid (Ruhmann et al, 2018), more degraded SFPs (Fricke et al, 2023), and higher levels of seminal fluid oxidative stress (Noguera et al, 2012) than ejaculates of young males. Thus, age-related changes in seminal fluid can play an important role in constraining the reproductive output of old males (Fricke et al, 2023; Sepil et al, 2020). In line with the disposable-soma theory of ageing (Maklakov and Immler, 2017), a higher rate of senescence in seminal fluid (SF) than sperm can be expected due to SF being produced by somatic tissue, which might be less protected from age-related damage than the germline (Fricke et al, 2023; Milholland et al, 2017). Measuring changes caused by both, gametic- and somatic-derived ejaculate components is crucial especially because each can senesce at different rates (Reinhardt et al, 2011; Sepil et al, 2020).

Male reproductive senescence might be particularly relevant in polygynous species, where males can mate with a series of multiple females in quick succession. Ejaculate production is costly (Dowling and Simmons, 2012; Olsson et al, 1997), and males in polygynous species can become depleted of sperm (Gerofotis et al, 2015; Pitnick and Markow, 1994; Rubolini et al, 2007) and seminal fluid (Hopkins et al 2019b; Linklater et al, 2007; Reinhardt et al, 2011; Sirot et al, 2009a) when mating-multiply. Such ejaculate limitation can decrease the reproductive output of polygynous males as they progress through series of successive matings (Abe, 2019; Lewis, 2004; Loyau et al, 2012; Macartney et al, 2020; Swierk et al, 2015). In such scenarios, the effects of male reproductive senescence might become more pronounced later in a mating sequence if old males suffer steeper rates of ejaculate depletion (Bressac et al, 2008, 2009, Appendix 1). For instance, terminal investment could lead old males to invest relatively more in early matings of a mating sequence, at the cost of lower investment in later matings (Duffield et al, 2017; Part et al, 1992), causing a steeper decline in reproductive output for old males through a mating sequence. Such hypotheses (see Appendix 1 for additional hypotheses) can only be tested by investigating age-specific patterns of sperm and seminal fluid depletion over a mating sequence. Surprisingly however, little is known about such patterns.

Here we use the fruit fly, *Drosophila melanogaster*, to investigate changes in the reproductive output of old and young males over a mating sequence, and subsequently investigate the ejaculate-mediated mechanisms underpinning these patterns. Fruit flies have been used extensively for reproductive ageing research. Previous studies show that old males have lower quality and quantity of sperm (Sepil et al, 2020; Turnell and Reinhardt, 2020) and seminal fluid (reviewed in Fricke et al, 2023), and lower reproductive success (Sanghvi et al, 2023; Snoke and Promislow, 2003) than young males. Male fruit flies are polygynous (Hopkins et al, 2019b), and due to a low sperm to egg number ratio (Bjork and Pitnick, 2006), can become ejaculate depleted after few matings (Pitnick and Markow, 1994; Hopkins et al, 2019b). However, the potential interacting effects of age and mate-multiplication on sperm and seminal fluid, and the fitness consequences of these interactions, have not been explored yet.

We combine sperm cell-labelling, biochemical and phenotypic approaches to investigate how ageing impacts reproductive output and ejaculate allocation strategies when mate-multiplying. We first test if advancing age leads to senescence in reproductive output of males in a mating sequence (Experiment A and Experiment B). Here, we additionally explore if old males increase investment in earlier matings compared to young males, in line with terminal investment hypothesis, or if old are more prudent in their ejaculate in early matings (Appendix 1). In experiment B, we then test if senescence in reproductive output can be mostly attributed to sperm limitation as is widely assumed. For this, we use fluorescence labelling and investigate how advancing age and mate-multiplying affect the number of sperm stored by males, transferred to and stored by females. Finally, we investigate whether senescence in seminal fluid (the somatic component of the ejaculate) might explain declines in male fertility with age (as predicted by the disposable soma theory). We test this by supplementing a female with extra seminal fluid from a young sterile male, prior to her mating with the young or old focal male, to test whether this partially rescues the declining fertility of old males (Experiment C).

## Results

### Experiment A

To compare the reproductive output (i.e. numbers of offspring produced) of old and young males over a mating sequence, we mated old or young wildtype *Dahomey* (*dah*) males to a maximum of 10 young, virgin *dah* females in quick succession over nine hours. The mated females were allowed to lay eggs for 24 hours to determine the number of offspring produced. Variation in the number of offspring produced by mated females was explained by a significant two-way interaction between male age and female number in a mating sequence (z = -2.24; P = 0.025; Figure 1A, Table S1). Post-hoc tests revealed that young males produced significantly more offspring compared to old males, earlier in a mating sequence. Overall, male reproductive output declined through a mating sequence, however this decline was steeper for young than old males (Figure S1).

**Figure 1:**
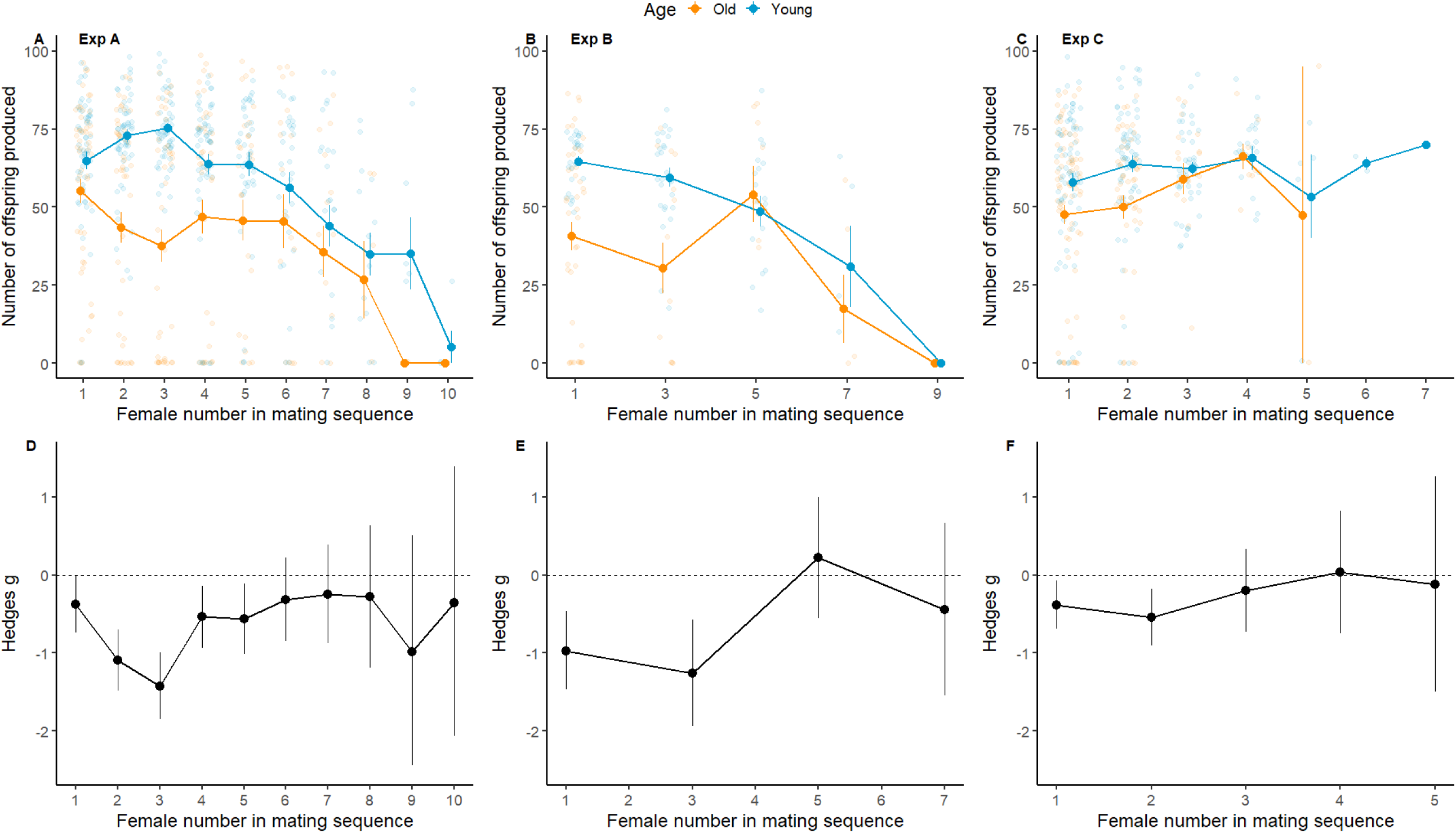
Reproductive output of old and young males in a mating sequence in experiments A, B and C. **1A, 1B, 1C**-Effect of male age and female number in a male’s mating sequence, on the number of offspring produced by mated females, for experiments (Exp) A, B, C respectively. Means and SE shown. **1D, 1E, 1F**-Post-hoc pair-wise comparisons between young and old males using effect sizes (Hedges’ g) of reproductive output with each female in a mating sequence, for experiments A, B, C respectively. Means and 95% C.I. shown. When C.I. does not overlap with zero, the effect is significant for α = 0.05. Negative values indicate old males having lower reproductive output than young males.

### Experiment B

Next, we wanted to test whether the lower reproductive output of old males seen in experiment A could be attributed to old males being sperm number limited, as commonly assumed. We used a transgenic line in which sperm heads are GFP-tagged (henceforth, “*gfp*” line), to count the sperm numbers produced by males, transferred to females and stored by females. However, we first ensured that the results seen in experiment A were consistent in experiment B. Like experiment A, we mated old or young *gfp* males to a maximum of 10 young, virgin *dah* females in a mating sequence over nine hours, to determine the number of offspring produced. There was no significant interaction between male age and female number in the mating sequence, on the number of offspring produced (z = -1.14, P = 0.254, Figure 1B, Table S2). However, variation in the number of offspring produced by individual females was explained by significant main-effects of male age (z = 2.01, P = 0.044) and female number (z = -3.23, P = 0.001). Specifically, old males produced fewer offspring than young males, and males produced fewer offspring with females later in a mating sequence (Figure 1B, 1E, Figure S2, Table S2). These results confirmed that age-dependent changes in offspring numbers were consistent across Experiment A and B.

Next, we tested whether this pattern was driven by differences in sperms numbers stored in old versus young males. For this, we compared the number of sperm present in the seminal vesicles (SV) of young and old *gfp* males. We found a significant interaction between male age and mating success (i.e. total number of females a male mated with), on the number of sperm present in a male’s SV (z = -4.63, P < 0.001, Figure 2A, Table S3).

**Figure 2:**
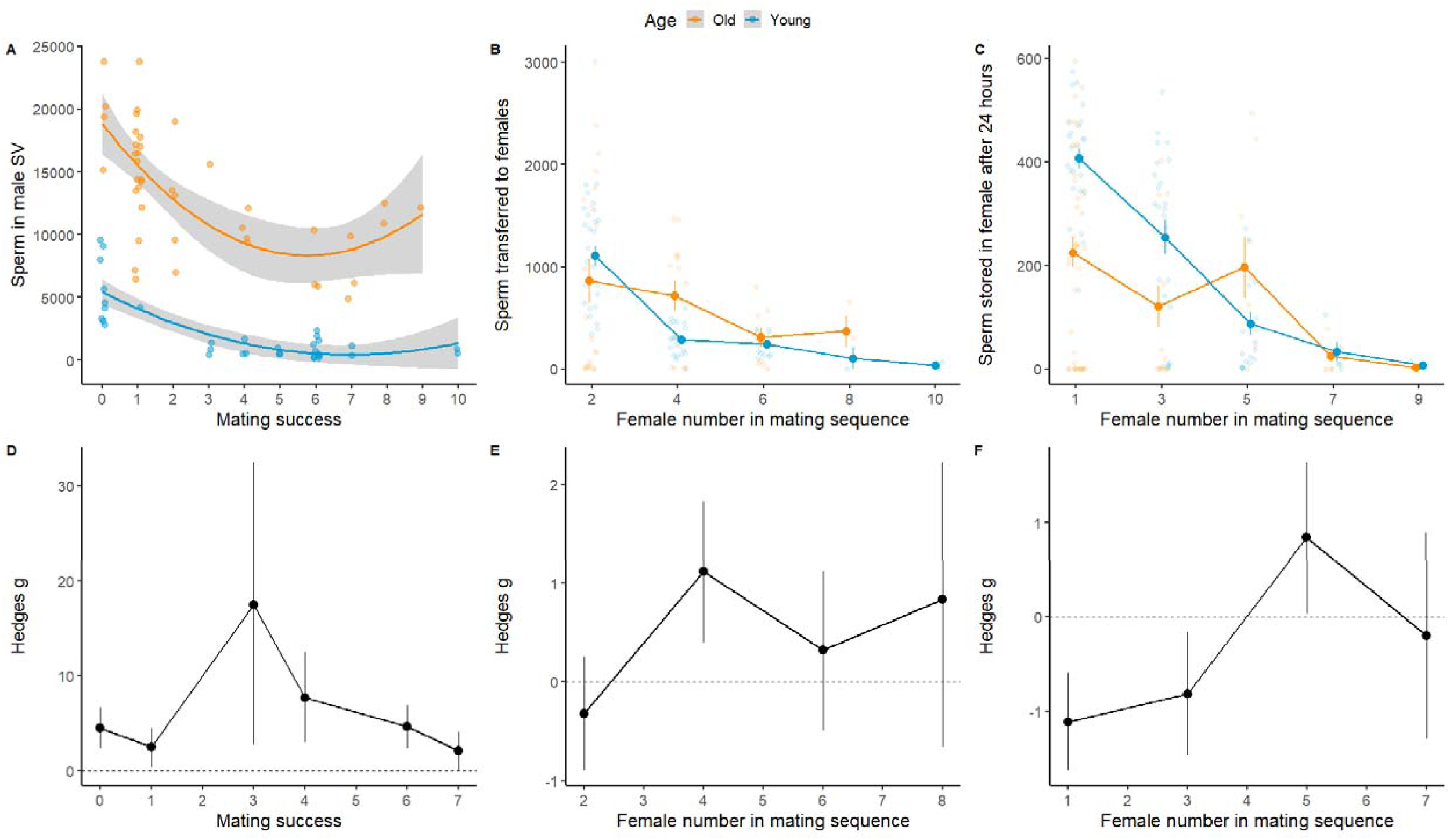
Age-dependent sperm accumulation and allocation through a mating sequence in experiment B. **2A-** Number of sperm in male seminal vesicles (SV) for old and young males with different mating success. **2B-** Sperm transferred to females by young and old males in a mating sequence. **2C-** Sperm stored by females mated to young and old males, after 24 hours. Means and SE shown for panels A, B, C. Panels **2D, 2E, 2F-** associated effect sizes when comparing young and old males in panels A, B, and C respectively, for each female number/mating success value. Means and C.I. shown for panels D, E, and F. For panels D, E, and F, when C.I. does not overlap with zero, the effect is significant for α = 0.05; negative values indicate that old males have lower sperm numbers than young males.

Specifically, old males had consistently more sperm present in their SV than young males, and this difference was greatest for males with intermediate mating success (Figure 2A, 2D). Hence, differences in offspring number could not be explained by older males being sperm limited.

We then explored whether the lower reproductive output of old males could be attributed to old males transferring fewer sperm to females. We found a significant two-way interaction between male age and female number in a mating sequence, to affect the number of sperm transferred to females (z = -2.905, P = 0.004, Table S4). Specifically, old and young males transferred similar numbers of sperm to females early in their mating sequence.

However, old males transferred more sperm to females later in the mating sequence in comparison to young males (Figure 2B, 2E, Figure S3). Therefore, age-related differences in offspring number could not be explained by older males transferring fewer sperm to females, compared to young males.

Next, we tested whether the low reproductive output of old males could be due to females storing fewer sperm when mated to old males, compared to young males. When comparing the number of sperm stored in a female’s long-term storage organs (seminal receptacle and spermathecae) 24 hours after mating, we found a significant two-way interaction between male age and female number in the sequence (z = -2.030, P = 0.042, Table S5). Specifically, early in a mating sequence, females mated to young males stored more sperm than females mated to old males (Figure 2C, 2F, Figure S4A). Visual inspection revealed an asymptotic relationship between the number of sperm stored by females after 24 hours, and the number of offspring produced by these females over 24 hours (Figure S4B). Generally, age-related changes in sperm stored in females was consistent with age-related changes in reproductive output of males, observed in Experiments A and B. Results on sperm numbers suggest that the low reproductive output of old males is likely due to females storing fewer sperm of old males.

In experiment B, we also indirectly tested whether accumulation and allocation of seminal fluid might explain the reproductive output phenotype observed in both experiments. For this, we compared the area of accessory glands (AG), which are the primary site of seminal fluid production in flies, of old and young *gfp* males. We found a significant interaction between male age and mating success (z = 6.577, P < 0.001, Table S6) on AG size. Old males had larger AG than young males, however the rate of decline in AG size was steeper for old than young males (Figure 3A). Therefore, later in a mating sequence, AG size did not differ between old and young males. Generally, old males had a higher ratio of sperm to AG size than virgin young males (Figure 3B), and this difference between old and young males became more exaggerated in males with higher mating success. These results suggest that old males might become seminal fluid depleted at a faster rate through a mating sequence than young males.

**Figure 3:**
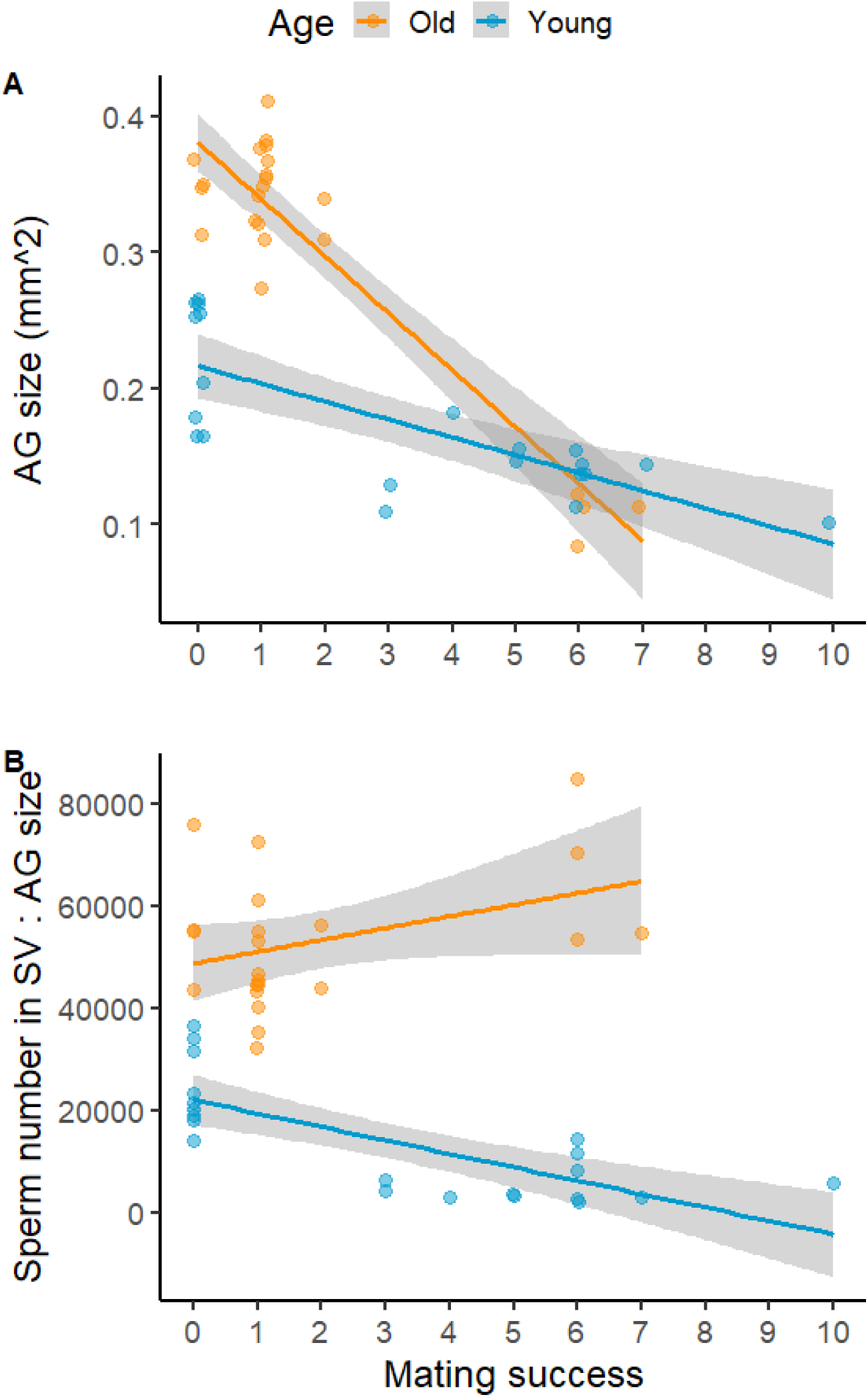
Relative and absolute changes in accessory gland size in old and young *gfp* males of different mating success in experiment B. **3A-** Significant interaction between male age and male mating success, to affect the size of male accessory glands (mm^2^). Old males, despite having larger AGs when virgins, show steeper declines in AG size than young males through a mating sequence. Means and 95% C.I. shown. **3B-** Ratio of sperm reserves in male SV to accessory gland size (count/mm^2^). Virgin old males have more sperm in their SV, than SF in their AGs, compared to virgin young males. Old males however allocate relatively more seminal fluid than sperm per mating while young males do the opposite, through a mating sequence. Young males were 3-7 days old, old males 37-42 days old.

### Experiment C

Finally, we tested if the decline in male reproductive output with age and over successive matings is driven by females being seminal fluid limited, given the documented role of seminal fluid in assisting sperm inside female for long-term storage. Specifically, we tested whether the receipt of “extra” young seminal fluid would rescue senescence in reproductive output seen in Experiments A and B. For this, we first mated young, virgin *dah* females to young, virgin *son of tudor* (“*sot”)* males (which are sterile, but transfer seminal fluid). These mated females that acquired “extra” seminal fluid from *sot* males, were then mated to the focal *dah* male (young or old) in a mating sequence (maximum of 10 matings over nine hours) on the same day, as in experiment A. After mating with *dah* males, the females were allowed to lay eggs for 24 hours to determine the number of offspring produced. We found no significant effect of male age (z = 1.65, P = 0.099), female number in a mating sequence (z = 1.3, P = 0.193), or an interaction between male age and female number to influence the number of offspring produced (z = -1.46, P = 0.145, Figure 1C, Figure S5; Table S7). When females previously received SF from young *sot* males, the reproductive output of focal old and young *dah* males was similar (Figure 1C, 1F), and male reproductive output did not decrease through a mating sequence. These results suggest that senescence in seminal fluid limits the reproductive output of old multiply-mating males.

In all three experiments, old males mated with fewer females (i.e. had lower mating success) compared to young males, which could be associated with longer mating latencies of old males (Appendix 2, Figure S6).

## Discussion

### Summary of findings

Male reproductive senescence has been demonstrated across taxa, in studies that mate males with an individual female (reviewed in Monaghan and Metcalfe, 2019; Sanghvi et al, 2024). However, these studies are not informative about reproductive senescence in polygynous species (Bressac et al, 2008, 2009), where males are exposed to multiple partners and male reproductive output can be modulated by female order in a male’s mating sequence (Abe, 2019; Lewis, 2004; Macartney et al, 2020; Appendix 1). Additionally, the pathways mediating male reproductive senescence can be diverse, and have been difficult to pinpoint. These include age-dependent declines in quality and quantity of sperm (Sanghvi et al, 2024) and seminal fluid (Fricke et al, 2023), changes in male behaviour and mating success (Dean et al, 2010; Hayes et al, 2013) and ejaculate allocation (Win et al, 2013). Furthermore, females might contribute to male reproductive senescence via preference for young males (Radwan, 2003; Vuarin et al, 2019). Our study bridges these knowledge gaps by investigating age-related changes in male reproductive output using multiple-mating assays, to understand the contribution of various processes causing male reproductive senescence.

Using three experiments and flies from three different lines, we investigated whether old and young males differed in their reproductive output through a mating sequence. We then explored whether observed patterns are explained by limitation of sperm numbers or seminal fluid. Overall, we found that old males had a lower reproductive output than young males. In general, male reproductive output declined through a mating sequence, and males produced fewer offspring with, and transferred fewer sperm to, females later in a mating sequence. However, this decline was steeper for young rather than old males (see Appendix 1 for predictions). We explored whether the lower reproductive output of old males was due to them being sperm depleted. Surprisingly, old males had larger sperm reserves, and transferred similar or greater numbers of sperm to females, than young males. However, females mated to old males stored fewer sperm compared to females mated to young males. We then explored whether seminal fluid might explain the lower reproductive output of old males. We discovered that when a female received extra seminal fluid (SF) from a young male prior to her focal mating, differences in reproductive output between old and young focal males diminished. This supplemented seminal fluid further alleviated declines in male reproductive output through a mating sequence. Old males had larger accessory glands (AGs), the main site of SF production and storage compared to young males. Yet, old males experienced steeper rates of decline in AG size through a mating sequence, relative to young males. Collectively, our results suggest that male reproductive senescence is less apparent later in a male’s mating sequence. Furthermore, our findings indicate that male reproductive senescence may be largely driven by senescence in seminal fluid or sperm quality, rather than senescence in sperm numbers.

### Changes with male age

We found that old males had higher numbers of sperm stored in their seminal vesicles than young males (also shown by Decanini et al, 2013; Kehl et al, 2013, 2015; Reinhardt et al, 2011). Sanghvi et al (2024) in a meta-analysis found a similar pattern across insects, and attribute this to experimental males typically being maintained as virgin, which can lead to longer periods of sperm accumulation in old than young males. Consistent with this, at low mating rates, old males often have larger ejaculates than young males, but at high mating rates this pattern is reversed (Aich et al, 2021; Bressac et al, 2009; Sepil et al, 2020). In our study, we maintained males as virgins until the mating assay, and old males might have accumulated more sperm than young males because spermatogenesis occurs throughout adult life in fruit flies with relatively low levels of sperm loss (Bjork et al, 2007; Demarco et al, 2013; Santos et al, 2023; Sepil et al, 2020). Old virgin males had a higher ratio of sperm reserves to AG size (thus SF reserves), compared to young virgin males, suggesting faster rates of sperm than SF accumulation, with age.

Despite having larger sperm reserves and transferring more or a similar number of sperm to females, females stored fewer sperm from old males (experiment B), and old males generally produced fewer offspring than young males (experiments A and B). Age-related changes in seminal fluid protein (SFP) composition or quality might explain our results. In experiment C, the prior receipt of seminal fluid (SF) by females resulted in similar number of offspring produced by old and young males. In fruit flies, SFPs produced in accessory glands play an important role in fertilisation, female sperm uptake and storage, and ovulation (Avila et al, 2011; Bloch Qazi and Wolfner, 2003; Chapman and Davies, 2004; Heifetz et al, 2005; Poiani, 2006; Ravi Ram and Wolfner, 2007). Old *D. melanogaster* males have lower expression of SFP genes such as ovulin, sex peptide and Acp36DE, and have more degraded SFPs compared to young males (Fricke et al, 2023; Koppik & Fricke, 2017; Sepil et al, 2020). Old *dah* males might have benefitted from females receiving seminal fluid from young *sot* males, explaining our results. Second, old males might have less viable or motile sperm than young males (e.g. Sepil et al, 2020; Sturup et al, 2013), leading fewer sperm to be stored by females and consequently reduce fertilisation rates for old males. Third, females mated to old males might have ejected greater proportions of sperm than females mated to young males, via cryptic female choice (Snook and Hosken, 2004; Vuarin et al, 2019; Wagner et al, 2004). Future experiments could use transgenic females that do not eject sperm and measure sperm viability, to test whether these might also explain our results.

Our finding of old males transferring similar sperm numbers as young males, but producing fewer offspring, might also be explained by variation in *quantities* of seminal fluid transferred. We found that old males had larger accessory glands, thus likely accumulated higher quantities of seminal fluid than young males, due to being kept as virgins (Reinhardt et al, 2011; Sepil et al, 2020). However, old fruit fly males, despite accumulating more seminal fluid and having larger AGs than young males, are known to transfer lower quantities of SFPs to females (Rezaei et al, 2015; Sepil et al, 2020). We suggest that future studies could manipulate the age of *sot* (SF-donor) males, as well as measure the quantity of SF transferred to females, to disentangle whether quality, composition, or quantity of SF explains our results better. Our results have important implications for sexual selection and conflict because they demonstrate that males who mate second might indirectly benefit from the SF of males who mate first with a female (Alonzo and Pizzari, 2010; Holman et al, 2009; Nguyen and Moehring, 2018). While not a direct test of the disposable-soma hypothesis (Maklakov and Immler, 2016), our results indicate higher rates of senescence in seminal fluid than sperm number, suggesting males might be investing more in maintaining the germline-derived than soma-derived components of the ejaculate.

Methodological factors might further explain why old males produced fewer offspring despite transferring similar or higher numbers of sperm to females. For instance, offspring from old fathers might have lower egg-to-adult survival (e.g. Preston et al, 2015; Ruhmann et al, 2018) due to deleterious paternal age effects (Monaghan et al, 2020). This could have biased our measurement of the number of eclosing offspring toward those of young males, even if old and young males fertilised equal numbers of eggs. Selective disappearance of males (Bouwhuis et al, 2009; Sanghvi et al, 2022) with low sperm production rates, especially in our *gfp* line that experienced high mortality (Figure S6), could have biased the cohort of old *gfp* males. Additionally, post-meiotic sperm damage (Pizzari et al, 2008; Wagner et al, 2004; White et al, 2008) could reduce the reproductive output of old males, due to old virgin males likely storing sperm for longer durations before ejaculation than young virgin males (Pizzari et al, 2008).

### Changes through a mating sequence

We found that males produced fewer offspring as they progressed through a mating sequence in Experiments A and B. Studies have previously shown that males, when multiply-mating, can become depleted of sperm and seminal fluid (Gerofotis et al, 2015; Pitnick and Markow, 1994; Rubolini et al, 2007; Sirot et al, 2009a), producing fewer offspring with females encountered later in a mating sequence (Abe, 2019; Lewis, 2004; Loyau et al, 2012; Macartney et al, 2020; Swierk et al, 2015). The asymptotic relationship between sperm stored in females, and offspring produced by females in our study, indicates that reproductive output through a mating sequence is likely limited by seminal fluid, not sperm numbers (Hopkins et al 2019b; Linklater et al, 2007; Reinhardt et al, 2011; Sirot et al, 2009a). Here, even if males transferred more sperm, offspring production would not increase because female ovulation and fertilisation rates are constrained by insufficient quantities of seminal fluid (Bloch Qazi et al, 2003; Chapman, 2001). Results from experiment C support this. Here, females were provided with SF from a previous mating before the focal mating took place, and focal male reproductive output did not decline through the mating sequence. Fruit fly males become depleted of seminal fluid at faster rates than sperm when multiply-mating (Reinhardt et al, 2011; Linklater et al, 2007), which is consistent with results of our study.

A lack of decline in reproductive output through a mating sequence in Experiment C might also be explained by male ejaculate plasticity. Females in this experiment were first mated to *sot* males before being used to mate with focal *dah* males. Male flies can detect female mating status using olfactory cues (Friberg, 2006; Lupold et al, 2011) to infer levels of sperm competition. Cues of high sperm competition risk (i.e. many rival males) lead male flies to transfer larger ejaculates to females (Bretman et al, 2009; Hopkins et al, 2019b; Price et al, 2012; Thomas and Simmons, 2009; Wedell and Cook, 1999) and even alter their SFP composition (Ramm et al, 2015; Wigby et al, 2009). Hence, in experiment C, focal *dah* males might have perceived higher sperm competition levels and increased their ejaculate investment in each mating. This could have alleviated a male’s reproductive decline through his mating sequence (possibly at the cost of fewer overall matings).

### Male age x mating sequence interaction

We discovered old males to have shallower declines in offspring production and sperm numbers transferred to and stored by females through a mating sequence, compared to young males. This result is unlikely due to prudent sperm allocation by old males (see Appendix 1) because old males had accumulated more sperm than young males. Our results do not support the terminal investment hypothesis either (Duffield et al, 2017; Froy et al, 2013), whereby we predicted terminal investment to lead old males to have steeper declines in reproductive output through a mating sequence, than young males. Instead, our results indicate that complex age-dependent sperm and seminal fluid accumulation and allocation patterns, lead to higher intercepts but steeper declines in reproductive output of young males through a mating sequence. Importantly, our results show that a female would benefit from mating with a young male only when she encounters him early in his mating sequence.

### Conclusions

Male reproductive senescence is multifaceted and can be either exacerbated or buffered under polygyny, depending on patterns of ejaculate accumulation and allocation, as well as the reproductive trait being measured. In polygynous species with low anisogamy such as *D. melanogaster* (Bjork and Pitnick, 2007), ejaculate depletion can have severe consequences for male and female reproductive and mating success. We show that different components of the ejaculate- sperm versus seminal fluid, decline at different rates with age and through a mating sequence. We show that old males have lower reproductive output than young males, which is likely driven by senescence in seminal fluid quality, sperm viability, or via female- mediated effects, rather than senescence in sperm number. These results have important consequences for sexual conflict and sexual selection (Adler et al, 2014; Damiens and Boivin, 2006; Carazo et al, 2011; see Appendix 3), whereby female fitness is not only determined by a male’s mating history (e.g. Wedell et al, 2002) or male age (Monaghan and Metcalfe, 2019), but also their interaction. Importantly, our results show that declines in male reproductive output are not permanent, and can be reversed by the presence of sufficient seminal fluid. This interplay between sperm and seminal fluid has crucial implications for our understanding of ageing and sexual selection, with possible biomedical implications.

## Methods

### Stock maintenance

All flies in our experiments were maintained at a 12:12hr light cycle at a constant temperature of 25°C and 45% relative humidity, under which, flies have an egg-to-adult developmental time of 10 days. We used males of the wildtype Dahomey strain (henceforth, “*dah”*) that has been maintained since the 1970’s in the lab. In experiment B, we additionally used *Ub-GFP* males (henceforth, “*gfp”*) where transgenes expressing green fluorescent proteins enable the visualization of sperm heads (Manier et al, 2010). In experiment C, we also used *son-of-tudor* (henceforth, “*sot”*) males that are infertile (i.e. sperm-less) but transfer seminal fluid (Boswell and Mahowald, 1985; Hopkins et al, 2019a; Kalb et al, 1993; Sepil et al, 2020; see Appendix 4). All females used in our experiments were young (3-4 days old) and of *dah* background. Across all experiments, male flies between 3 to 11 days of age were considered young, and between 37 to 46 days as old, based on previous studies (Aguilar et al, 2023; Sanghvi et al, 2023; Sepil et al, 2020; Snoke and Promislow, 2003; Ruhmann et al, 2018; Turnell and Reinhardt, 2020).

### Experimental flies

To generate experimental males of each line (*dah*, *gfp,* and *sot*), we collected eggs from our stock populations with a standardised egg density of ∼150 flies per bottle (Clancy and Kennington, 2001). To ensure virginity, we collected flies within six hours of emergence, using ice anaesthesia (Clancy and Kennington, 2001). All experimental males and females were fed with Lewis medium supplemented with molasses and *ad libitum* live yeast (Lewis, 1960). All experimental flies were kept as virgins in single-sex vials of 10 individuals, until being used for the experimental assays (more below). Experimental virgin males were transferred onto new food once a week. Virgin males in our lab usually have a median and maximum lifespan of 50 and 90 days respectively (Sanghvi et al, 2023; Sepil et al 2020; also see Figure S7). Sample sizes across the three experiments are provided in Table S8.

### Mating experiments

<u>Experiment A-</u> We first compared changes in the reproductive output of young (9-10 days old) and old (44-46 days old) males in a multiple-mating sequence. On the day of the mating assay, 60 old and 60 young experimental *dah* males were haphazardly chosen to successively mate with a maximum of *dah* 10 females, over a duration of 9 hours. Each male was moved into a vial containing a single young virgin female and observed. Each male remained with a female until the pair mated, and if a male did not mate with a female by the end of the 9 hour assay, that female was not included in our analysis. Once mated, the male was immediately transferred into another vial containing a new, young virgin female. Mated females were given 24 hours to oviposit in the same vial, after which females were discarded. The flies (i.e. offspring) in these vials were given 14 days to develop and eclose, after which these vials were frozen at -20°C and the number of eclosed (adult) offspring counted. This experiment was conducted across two replicates (blocks).

### Experiment B

Next, we investigated whether sperm limitation might drive differences in reproductive output of old and young mate-multiplying males. Specifically, we compared the number of sperm transferred to, and stored by, females mated to old and young mate- multiplying males, as well as sperm reserves and accessory gland size of mate-multiplying males. For this, we first generated experimental old (37 to 41 days old) and young (3 to 7 days old) *gfp* males. We then haphazardly chose 50 old and 28 young males to successively mate with a maximum of 10 young virgin *dah* females, over a duration of 9 hours. Old males are less likely to mate thus more were used. Like experiment A, each male was moved into a vial containing a single female and following a mating, males were immediately transferred into a new vial containing a different female. Following mating, odd-numbered females in a male’s mating sequence (i.e. 1^st^, 3^rd^, 5^th^, 7^th^, and 9^th^ female), were given 24 hours to oviposit in the same vial, after which the female was frozen at -20°C. These vials were frozen 14 days after oviposition, and the number of eclosed (adult) offspring in these vials were counted. All even-numbered females in a male’s mating sequence (i.e. the 2^nd^, 4^th^, 6^th^, 8^th^, and 10^th^ female) were frozen within 30 minutes of mating, at -20°C. Additionally, all mated *gfp* males as well as four old and nine young virgin *gfp* males not exposed to females, were frozen at -20°C after the mating assay. Reproductive tracts of frozen females (bursa, seminal receptacle, and spermathecae) and males (seminal vesicle and accessory glands) were dissected in PBS under a Leica M80 dissection microscope (Appendix 5, 6, 7). Male accessory glands (the primary site of seminal fluid production in flies) and the sperm stored in odd-numbered females were imaged using a Nikon E600 fluorescence microscope (Appendix 7). Sperm stored in even-numbered females and in male seminal vesicles were imaged using a Zeiss LSM880 confocal microscope (Appendix 5, 6). Data from images was obtained using FIJI (Schindelin et al, 2012; Appendix 5, 6, 7). This experiment took place across three replicates (blocks).

### Experiment C

Finally, we tested whether seminal fluid limitation might explain differences in the reproductive output of old and young, multiply-mate males. To test this, we investigated whether seminal fluid obtained by a female from her first mating (with a young *sot* male), impacts the reproductive output of old and young *dah* males, who subsequently mate with these females. For this, we generated young (4 to 11 days old) virgin *sot* males which lack sperm but produce seminal fluid (see Appendix 4). We mated ∼200 young (3 to 4 days old) virgin *dah* females, each with a young virgin *sot* male, and observed their matings. On the same day, we then conducted a multiple-mating assay using these *sot*-mated *dah* females, and experimental young (9-10 days old) and old (44-46 days old) *dah* males as previously described in experiment A. Specifically, we chose 115 old and 80 young experimental *dah* males haphazardly to successively mate with a up to 10 *sot*-mated *dah* females, over a duration of 9 hours. More old males were used because old males were less likely to mate with females. Each young or old experimental *dah* male was moved into a vial containing a single *sot*-mated *dah* female, and observed. Once mated, experimental *dah* males were immediately transferred into a new vial containing a different, young *sot*-mated *dah* female. Females who mated with experimental *dah* males were given 24 hours to oviposit in the same vial, after which females were discarded. These vials were frozen 14 days after the oviposition period at -20°C, and the number of eclosed (adult) offspring in the vials were later counted. This experiment was also conducted across two replicates (blocks).

### Data analysis

#### Male reproductive output

##### Experiments A, B, C

We analysed data on male reproductive output through a mating sequence from each of the three experiments separately, using the package *glmmTMB* (Brooks et al, 2017) in R v4.2 (R core team, 2012). In each of these analyses, the numbers of offspring produced (i.e. reproductive output) by each female over 24 hours of oviposition, was our dependent variable. We included male age (young or old), female order (henceforth, female number) in a mating sequence (1-10), their two-way interaction (i.e. male age x female number), and replicate, as fixed effects, with male ID as a random effect across all models. Additionally, we included observation-level (i.e. row) as a random effect to control for overdispersion in the data (Harrison, 2014), which was assessed using *DHARMa* (Hartig, 2017). Each analysis involved two steps. First, we ascertained whether including a zero-inflation term improved model fit. We did this by comparing a model with Poisson against one with a zero-inflated Poisson error distribution, and chose the model with the lowest AIC (henceforth, best-fit distribution). Second, using our best-fit distribution model, we compared three different fixed effect structures pertaining to male age and female number, to understand whether changes in male reproductive output though a mating sequence was linear or not. These model comparisons were done using a likelihood ratio test with the function *anova* in the package *base* (R core team, 2012). These fixed effect structures were: a model where male age interacted with only the linear term of female number (age * female number); one where male age interacted separately with both, the linear and quadratic term for female number ((age * female number) + (age * I(female number^2))); and one with male age, a linear term for female number, their two-way interaction, and a separate quadratic term for female number ((age * female number) + I(female number^2)). Importantly, males that did not copulate with a single female were excluded from these analyses.

### Sperm transfer and storage

#### Experiment B

We compared the number of sperm in the seminal vesicles of old and young males in experiment B. Here, we included male age, male mating success (i.e. total number of females a male mated with), their two-way interaction, and replicate, as fixed effects. We modelled sperm counts (estimated using the ‘find maxima’ plugin on FIJI, Appendix 7) as our dependent variable (one value per male). Similar to the models on reproductive output described above, for our model on sperm numbers in males, we first determined the best-fit error distribution. We then determined the most suitable fixed effects terms pertaining to male mating success and whether to include them as a linear or quadratic term.

We additionally analysed data on the number of sperm transferred to odd-numbered females, and the number of sperm stored by even-numbered females after 24 hours of egg laying, in two separate models. To test how male age affected the number of sperm transferred to odd-numbered females, we modelled sperm number as our dependent variable. Here, sperm numbers were estimated automatically from images of female bursa, seminal receptacle, and spermathecae, using the ‘find maxima’ plugin on FIJI win32 (see Appendix 6). To test how male age affected the number of sperm stored by even-numbered females, we modelled sperm number as our dependent variable. Here, sperm numbers in female spermathecae and seminal receptacle were counted manually using the cell counter plugin on FIJI win32 (see Appendix 6). For both models on sperm number in females, we modelled male age, female number in a mating sequence, their two-way interaction, and replicate, as fixed effects, with male ID and an observation-level as random effects. Our model selection procedure for both models was the same as described above (see data analysis: male reproductive output). Specifically, we first determined the best-fit error distribution for our models, and then determined the best fixed effects pertaining to female number being linear or quadratic.

Lastly, we analysed data on the area of accessory glands (which are the primary site of SF production) on a subset of males. For this, we created a linear model with Gaussian error distribution in the *lme4* (Bates et al, 2014) package, and included male accessory gland area (one value per male, which was the average area of the two accessory glands) as our dependent variable. Male age, male mating success (i.e. total number of females a male mated with), their two-way interaction, and replicate, were modelled as fixed effects. All LMMs were checked for normality of residuals and homoscedasticity, using the *stats* package (R core team, 2012).

### General notes on analysis

For all our models, female number was included as an ordinal variable. Generally, only when the two-way interactions between male age and female number or male mating success were non-significant, we created a main-effects model to interpret the independent influence of fixed effects (Engqvist, 2005). For all our models, we conducted post-hoc pairwise comparisons between old and young males using effect sizes (Hedge’s g, with α = 0.05) for each female number in a mating sequence. Final model structures and the variance explained by fixed effects in final models (R^2^_marginal_) are described in Table S9.

## Data and code availability

All data from our experiment and the R code used for analysis can be found at OSF: 10.17605/OSF.IO/5Z7M3. Images of dissected *gfp* males’ SV, AGs, and images for sperm numbers transferred to and stored by females, can be found on figshare: <u>10.6084/m9.figshare.25407223</u>, <u>10.6084/m9.figshare.25407298</u>, <u>10.6084/m9.figshare.25407253</u>, <u>10.6084/m9.figshare.25407382</u>.

## Supplementary figures

**Fig S1:**
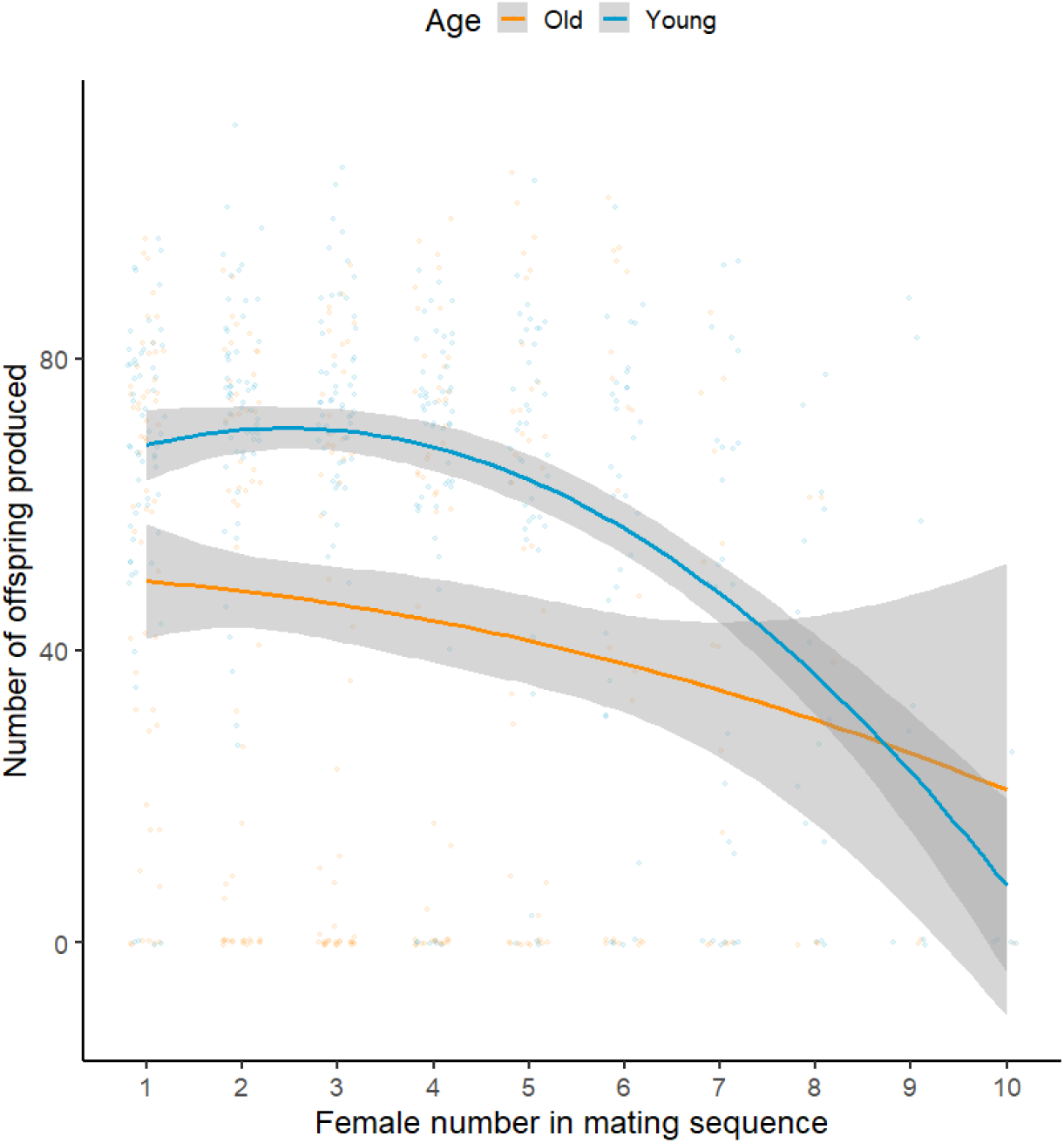
Significant interaction between male age and female number in a male’s mating sequence, to affect the number of offspring produced by a female over 24 hours of egg laying, in Experiment A. Old males produce fewer offspring than young males only early on in a mating sequence. Means and 95% C.I. shown.

**Fig S2:**
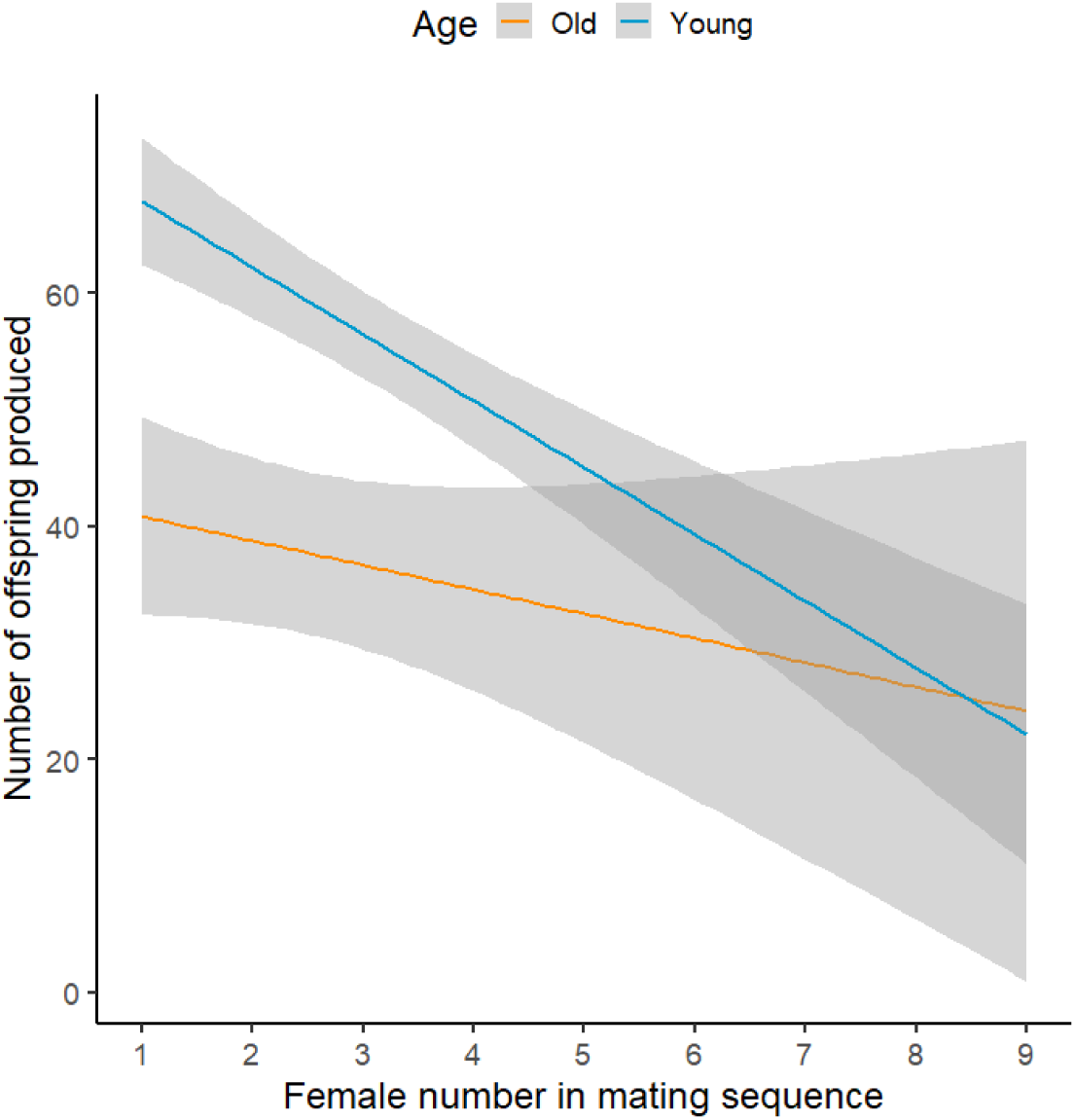
Significant effect of male age and female number in a male’s mating sequence, to affect the number of offspring produced by a female over 24 hours of egg laying, in Experiment B. Old males produce fewer offspring than young males, and males produce fewer offspring with females later than earlier in a mating sequence. Means and 95% C.I. shown.

**Figure S3:**
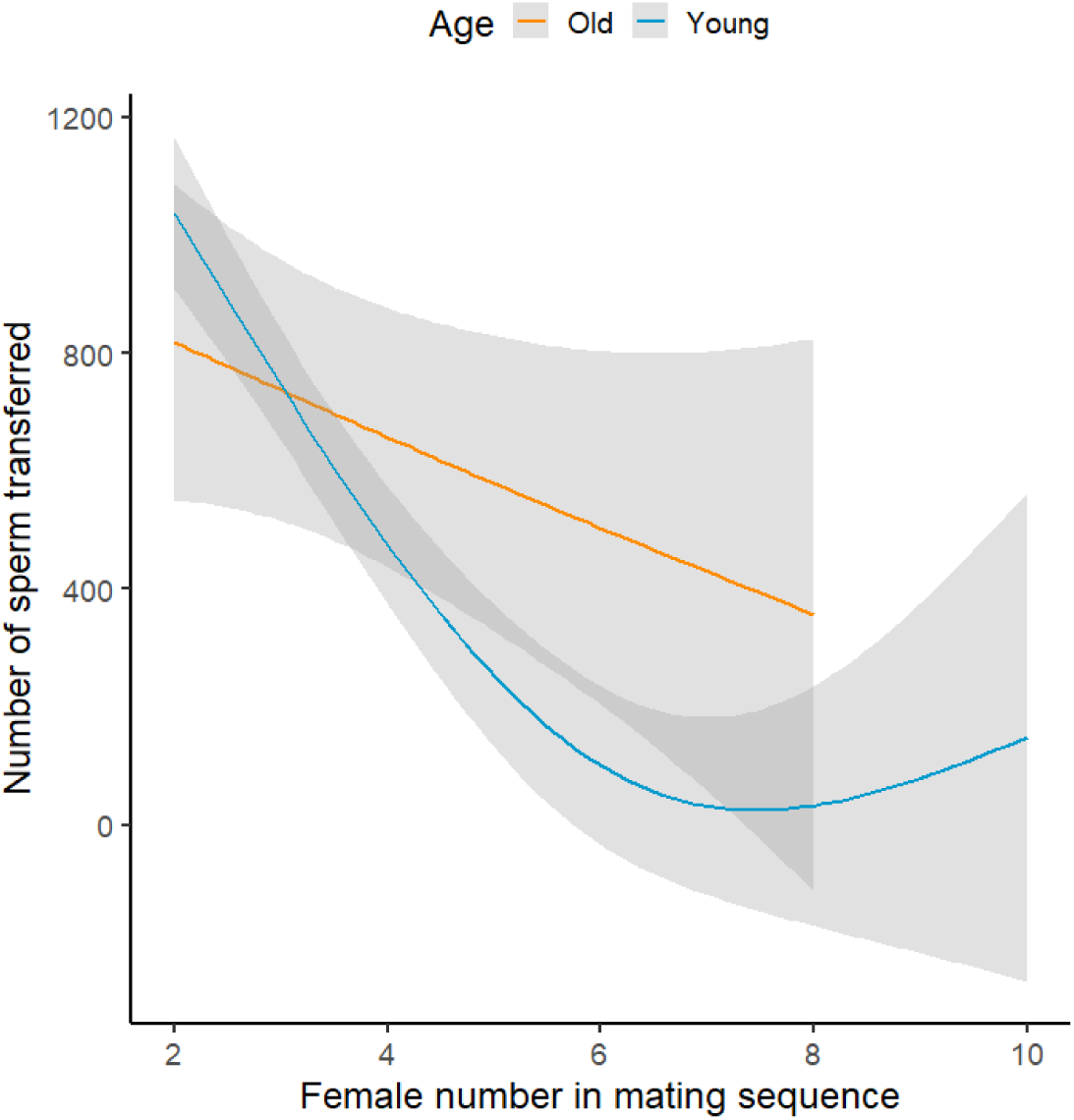
Effect of male age and female number in a male’s mating sequence, on the number of sperm transferred by the male to the female, in experiment B. Means and 95% C.I. shown.

**Figure S4:**
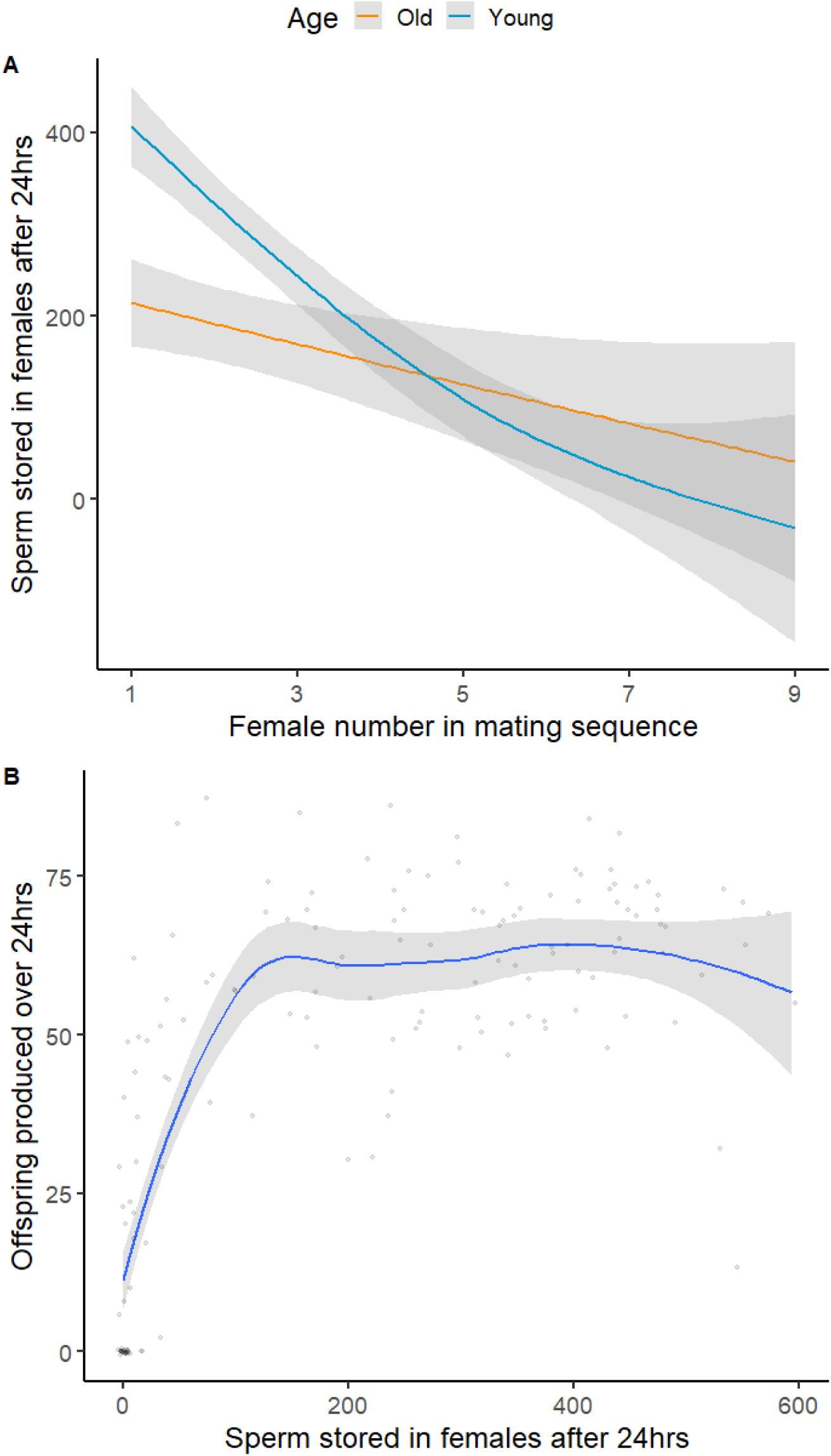
Experiment B. 4A-Effect of male age and female number in a male’s mating sequence, on the number of sperm stored by mated females after 24 hours of egg laying. 4B- Co-variance between number of sperm stored in odd-numbered females after 24 hours and the number of offspring produced by these females over 24 hours. Plot created using a loess smooth, to illustrate the non-linear (asymptotic) relationship. Means and 95% C.I. shown.

**Fig S5:**
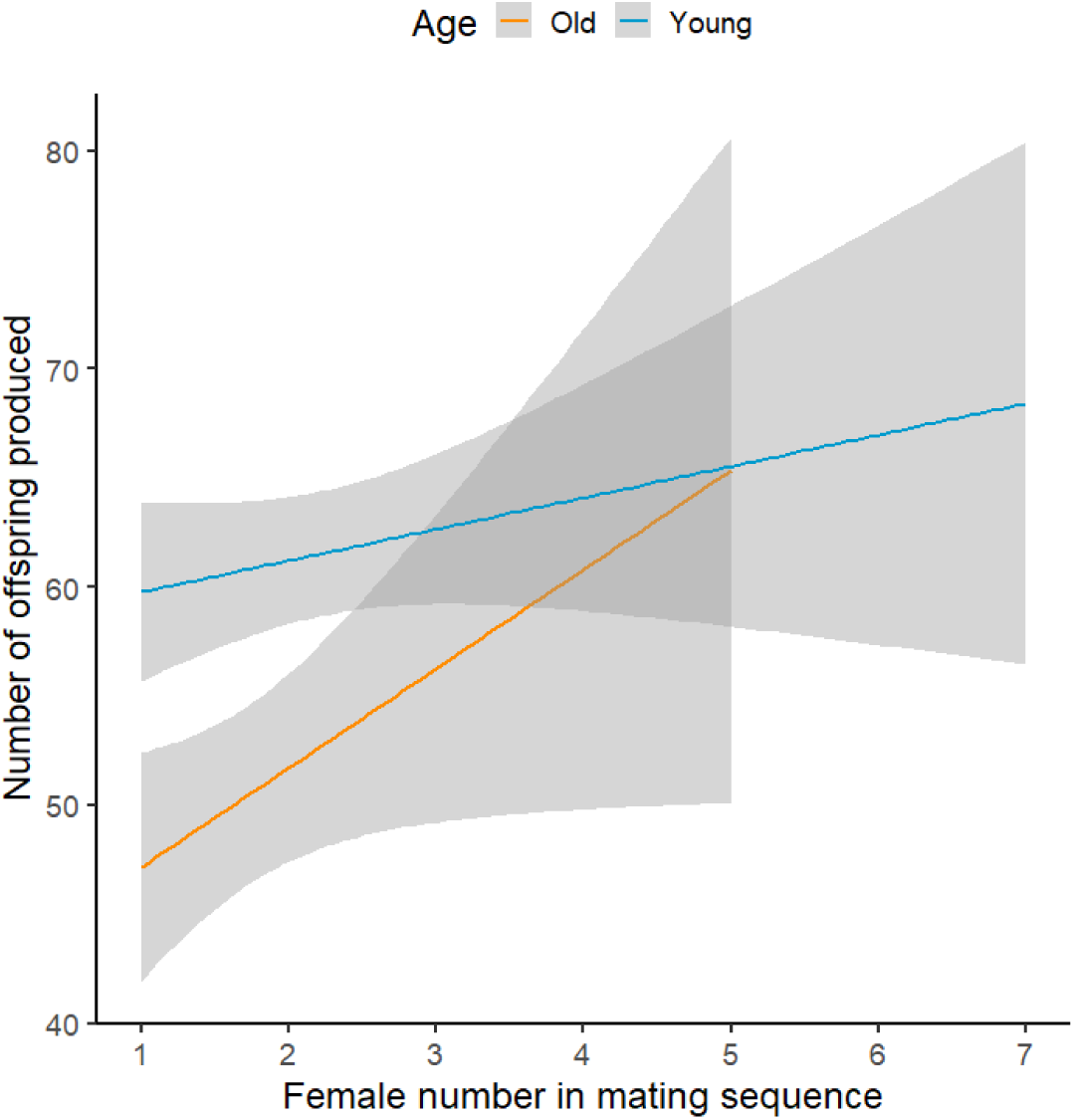
No significant effect of male age or female number in a male’s mating sequence, to affect the number of offspring produced by a female over 24 hours of egg laying, in Experiment C. Females, prior to focal mating with old or young *dah* males, were first mated with *sot* males to provide females with “extra” seminal fluid. Means and 95% C.I. shown.

**Figure S6:**
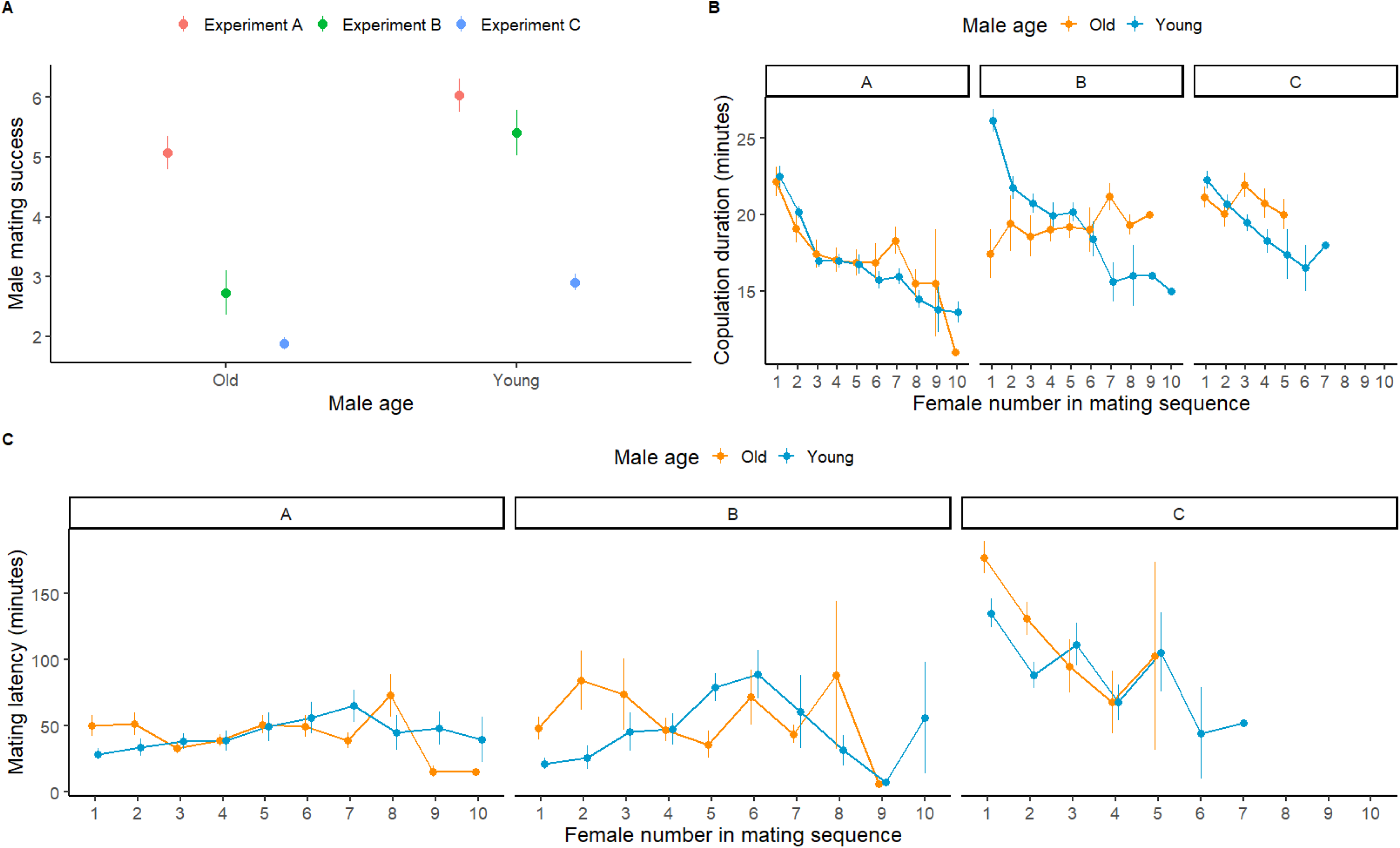
**A**. Mating success of focal old and young males used in our three experiments. **B.** Copulation duration of focal old and young males across Experiments A-C. **C.** Mating latency of focal young and old males across experiments A, B, and C. Means and SE shown.

**Figure S7:**
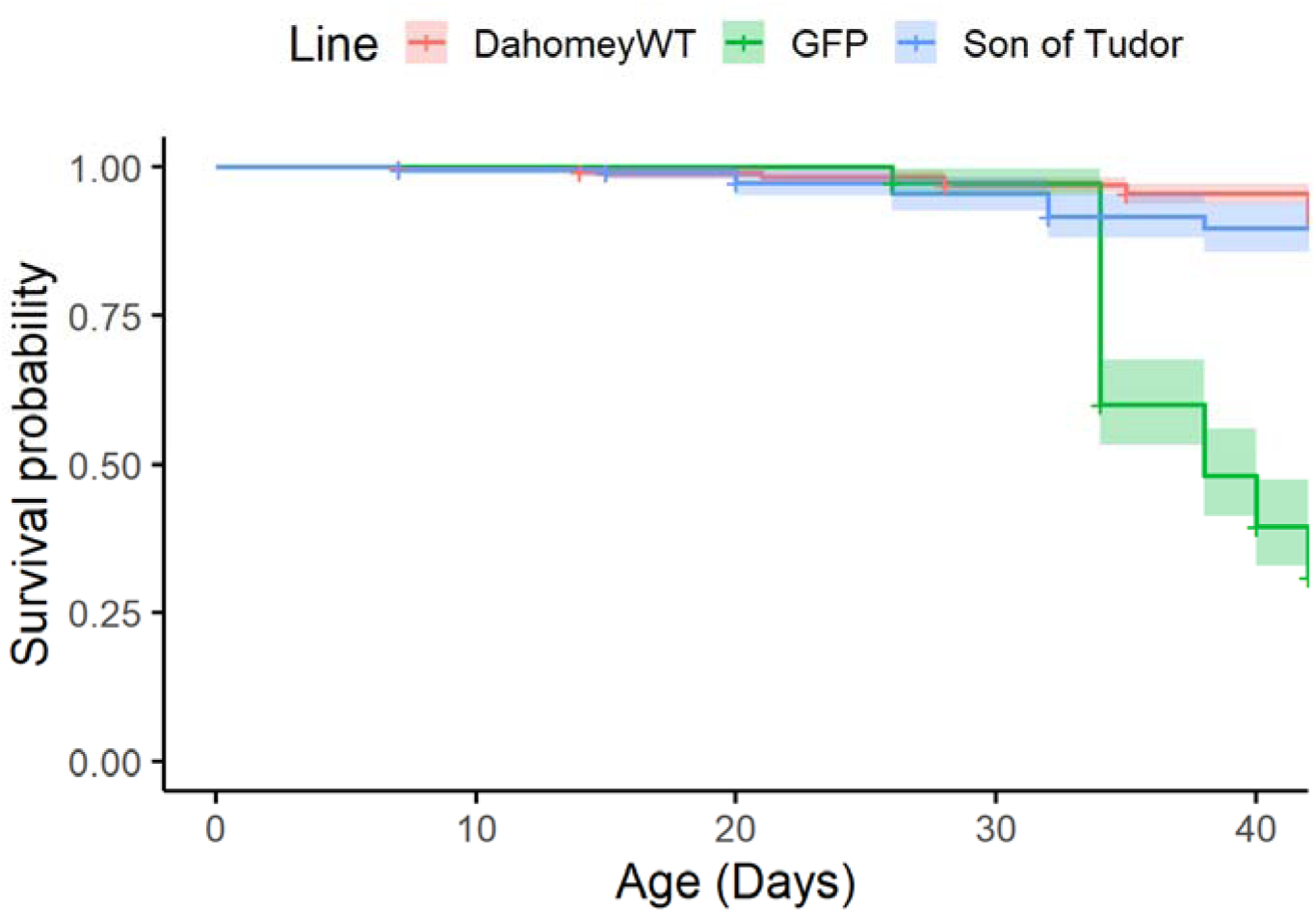
Age-dependent survival probability of males used in our study, from three distinct lines of *Drosophila melanogaster: dah, gfp, and sot. Gfp* males experienced higher mortality than *sot* or *dah* males. Means and 95% C.I. shown.

## Supplementary Tables

**Table S1:**
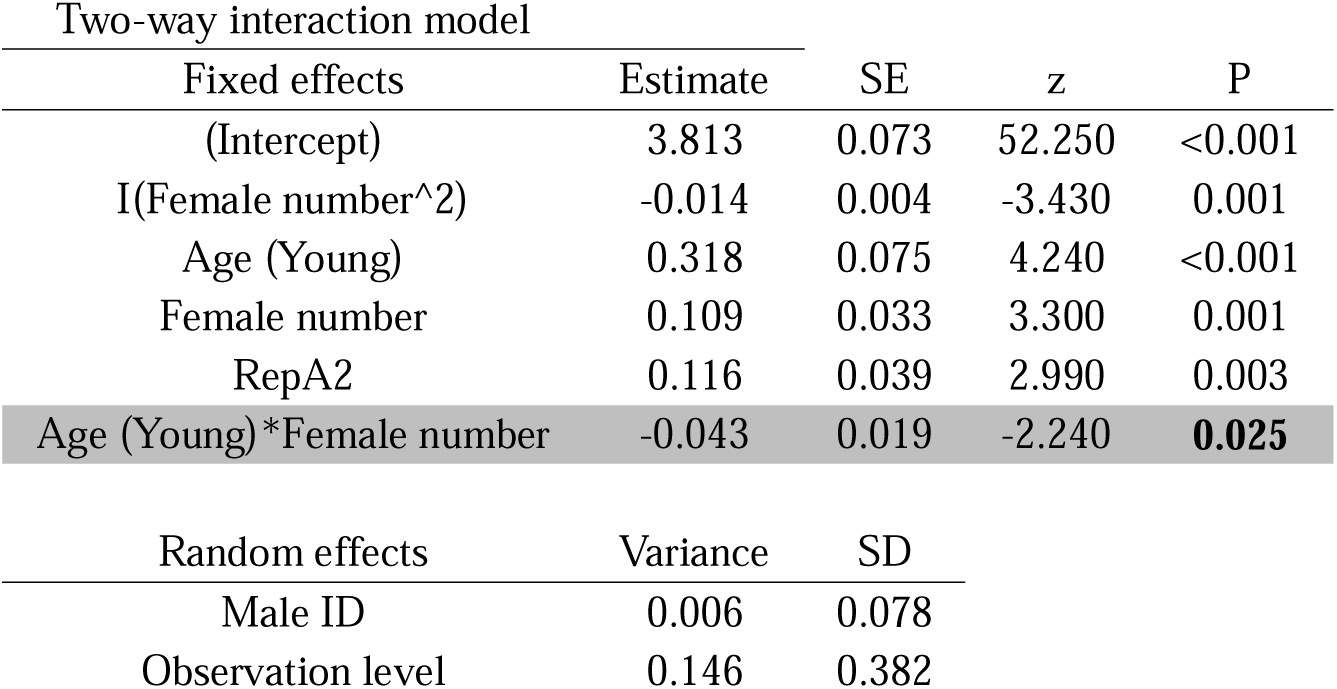
Effects of male age and female number in a male’s mating sequence, on the number of offspring produced by mated females in Experiment A. Model constructed with zero inflated Poisson error distribution. Two-way interaction model used to interpret interaction only, main-effects model used to interpret main-effects only when two-way interaction is non-significant. Effects of interest highlighted in grey, significant P values of interest in bold.

**Table S2:**
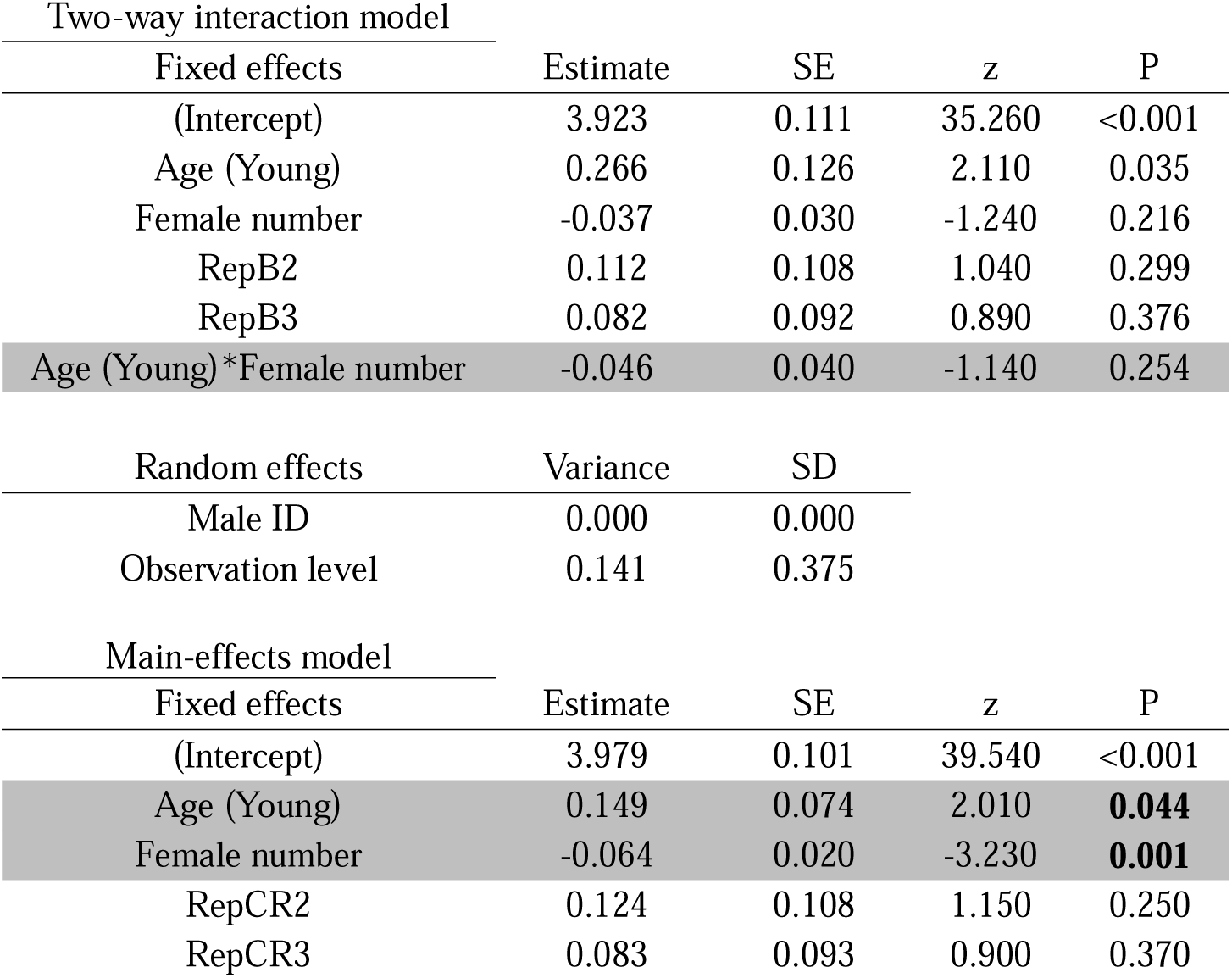
Effects of male age and female number in a male’s mating sequence, on the number of offspring produced by mated females in Experiment B. Model constructed with zero inflated Poisson error distribution. Two-way interaction model used to interpret interaction only, main-effects model used to interpret main-effects only when two-way interaction is non-significant. Effects of interest highlighted in grey, significant P values of interest in bold.

**Table S3:**
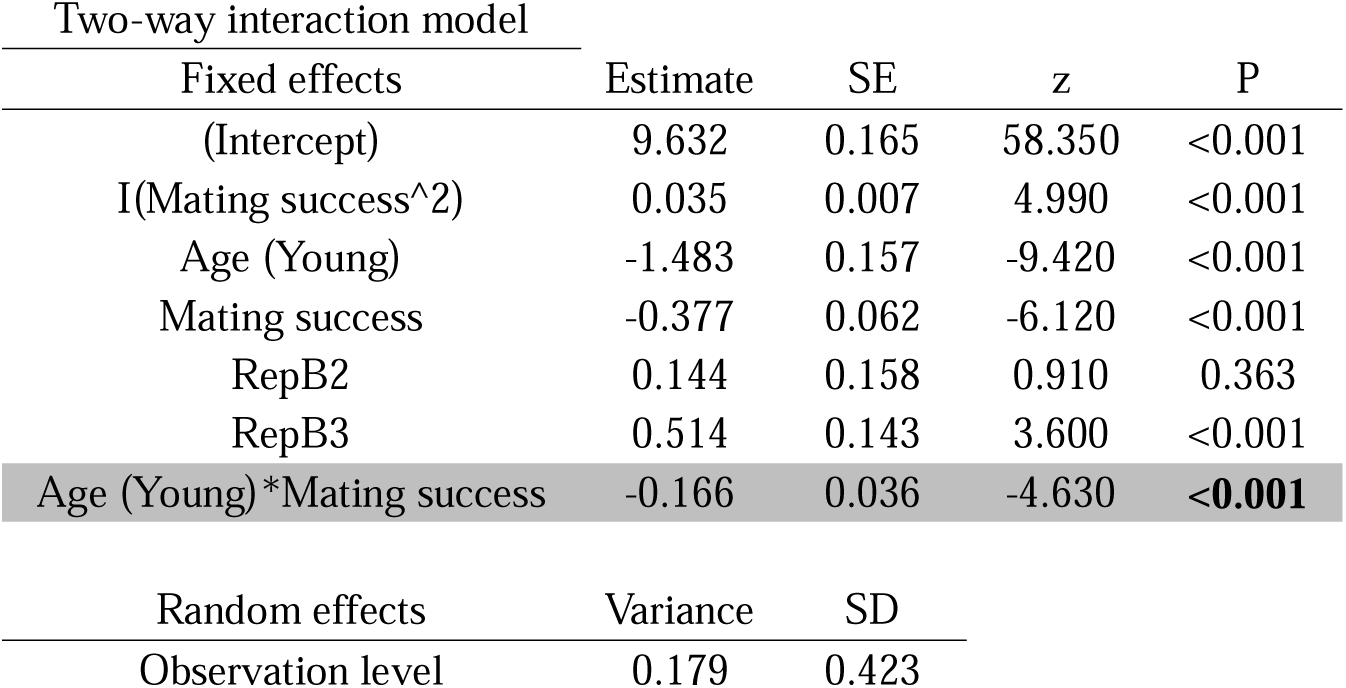
Effects of male age and male mating success, on the number of sperm accumulated in seminal vesicles of males in experiment B (one data point per male). Model constructed with Poisson error distribution. Two-way interaction model used to interpret interaction only, main-effects model used to interpret main-effects only when two-way interaction is non-significant. Effects used for interpretation highlighted in grey, significant P values of interest in bold.

**Table S4:**
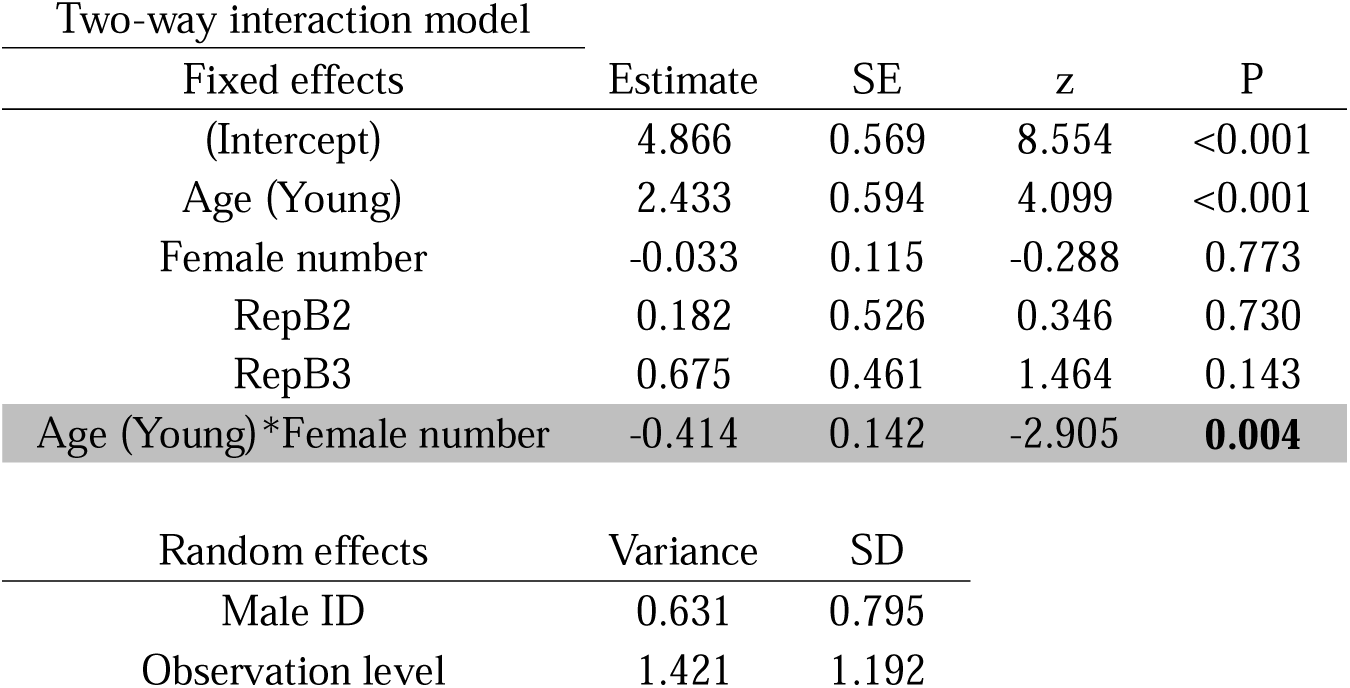
Effects of male age and female number in a male’s mating sequence, on the number of sperm transferred by males to mated females in Experiment B. Model constructed with zero inflated Poisson error distribution. Two-way interaction model used to interpret interaction only, main-effects model used to interpret main-effects only when two-way interaction is non-significant. Effects used for interpretation highlighted in grey, significant P values of interest in bold.

**Table S5:**
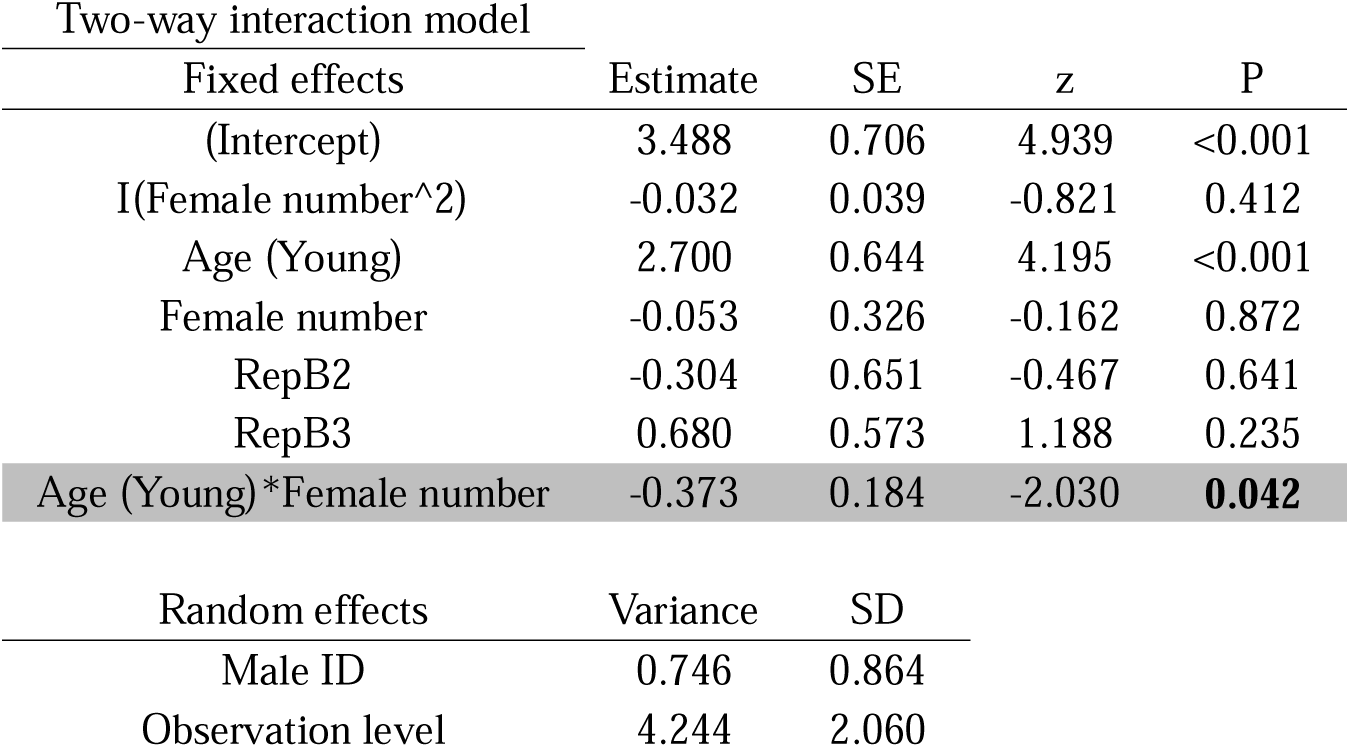
Effects of male age and female number in a male’s mating sequence, on the number of sperm transferred stored by mated females in Experiment B, after 24 hours of egg laying. Model constructed with Poisson error distribution. Two-way interaction model used to interpret interaction only, main-effects model used to interpret main-effects only when two-way interaction is non-significant. Effects used for interpretation highlighted in grey, significant P values of interest in bold.

**Table S6:**
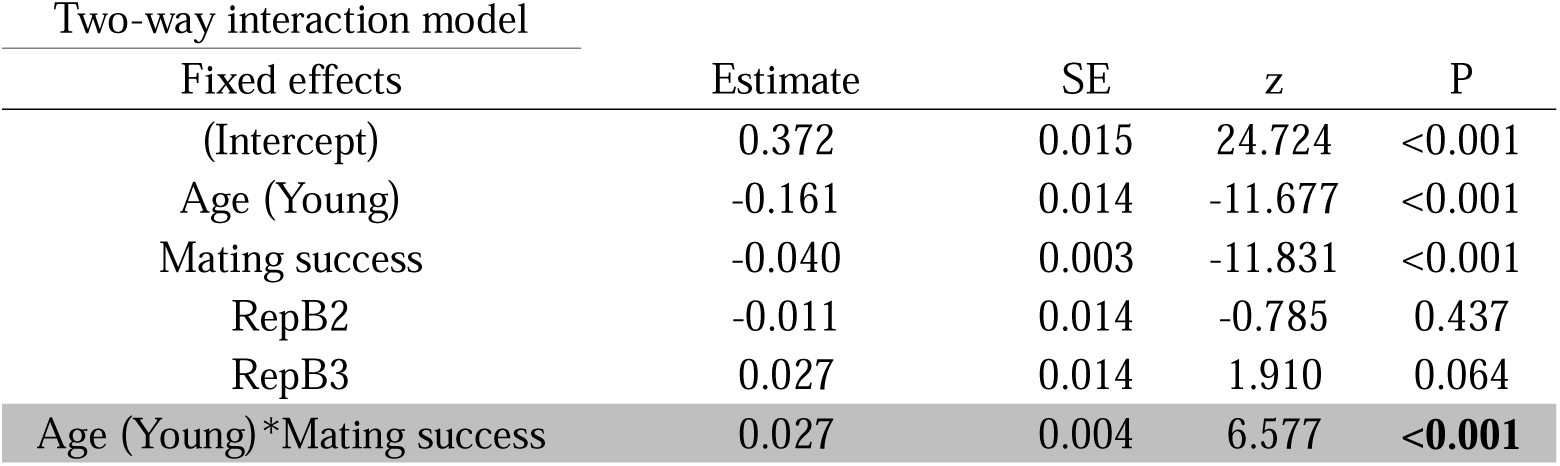
Effects of male age and male mating success, on the area of male accessory glands (cm^2^) in experiment B (one data point per male). Model constructed with Gaussian error distribution. Two-way interaction model used to interpret interaction only. Effects used for interpretation highlighted in grey, significant P of interest values in bold.

**Table S7:**
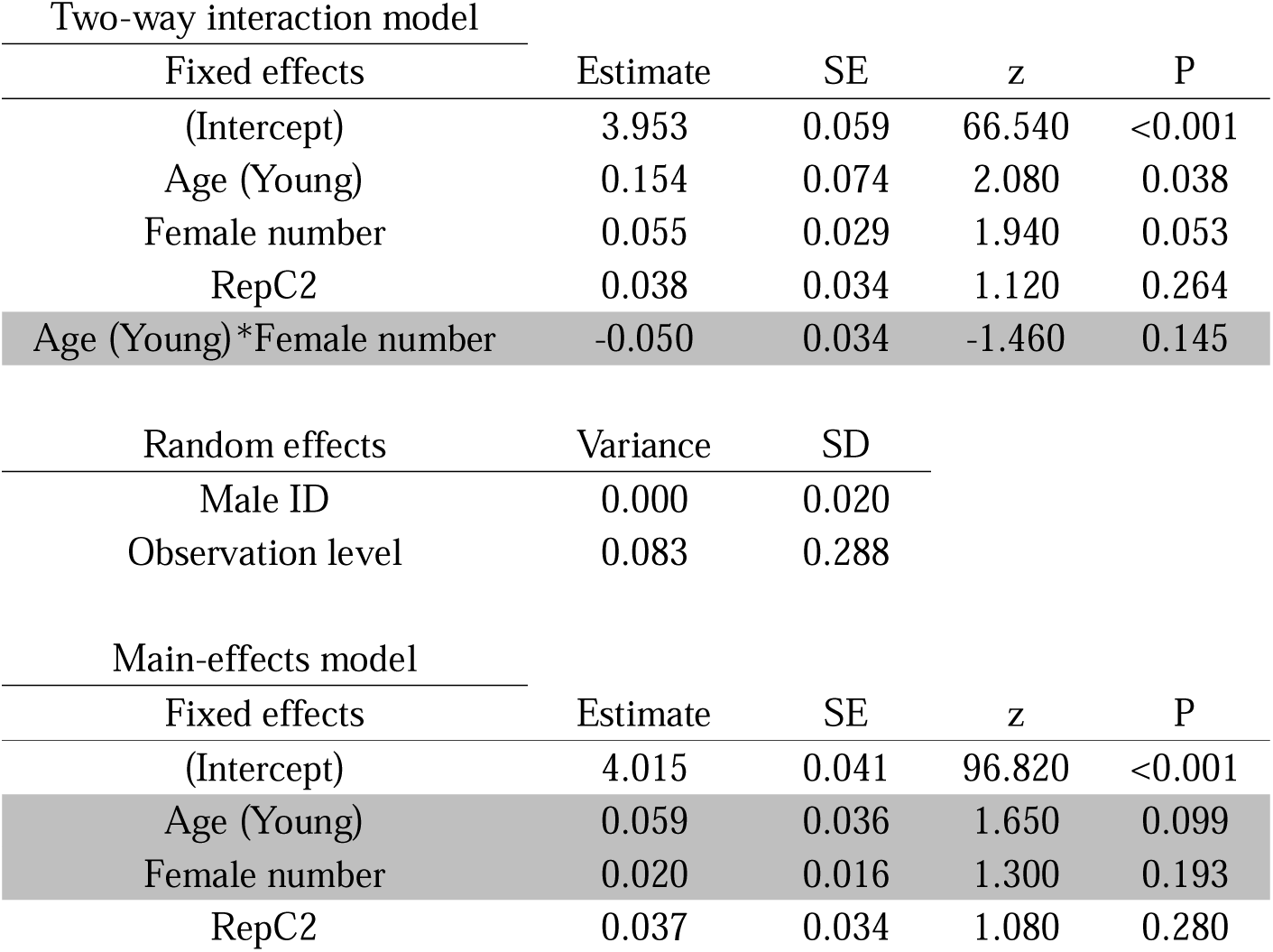
Effects of male age and female number in a male’s mating sequence, on the number of offspring produced by mated females in Experiment C. Model constructed with zero inflated Poisson error distribution. Two-way interaction model used to interpret interaction only, main-effects model used to interpret main-effects only when two-way interaction is non-significant. Effects of interest highlighted in grey, significant P values of interest in bold.

**Table S8a:**
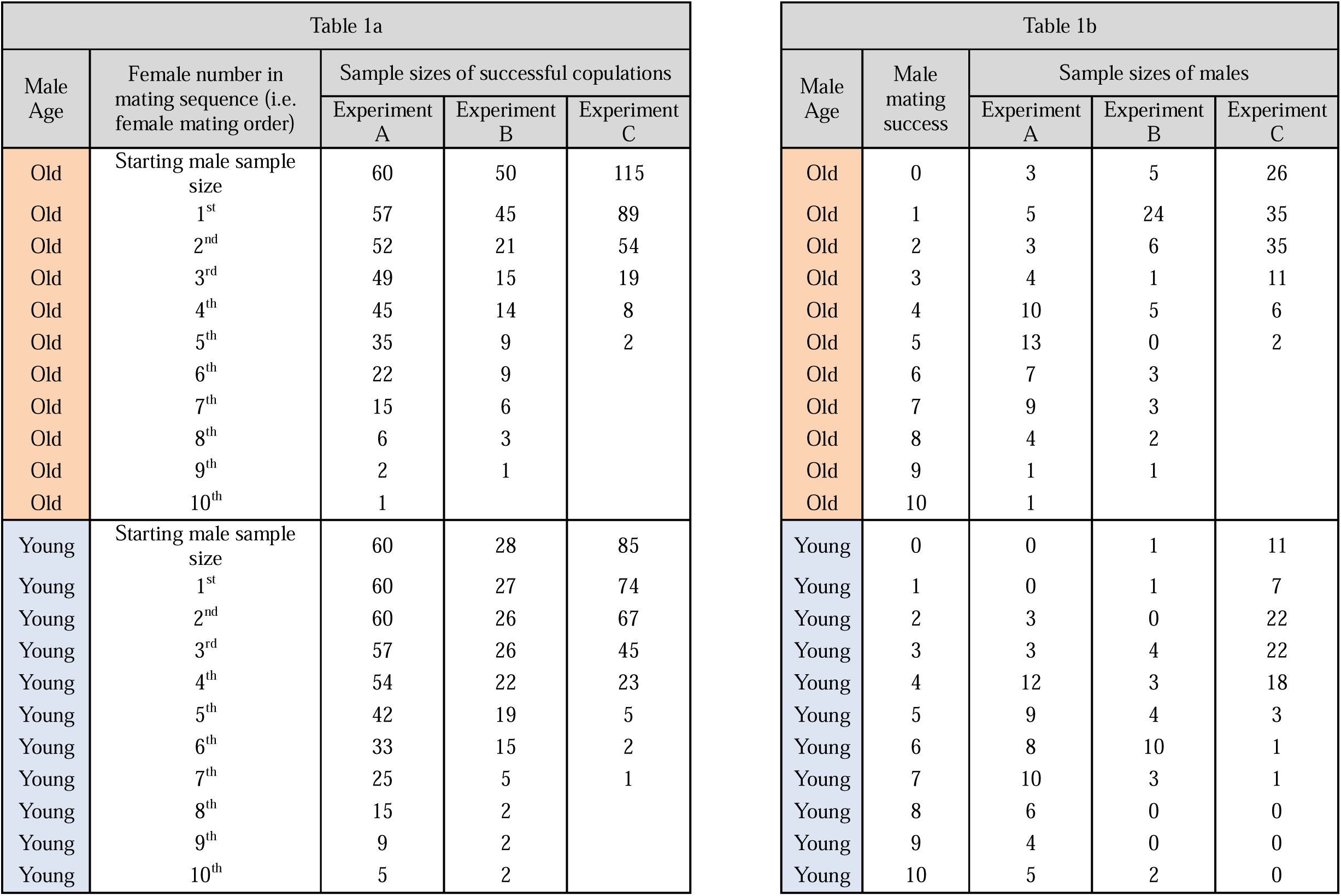
sample sizes for number of successful copulations by old and young males with females in a mating sequence, for Experiments A-C. Table 1b: sample sizes for total mating success (i.e. sum of females a male mated with) of old and young males, in Experiments A-C.

**Table S9:**
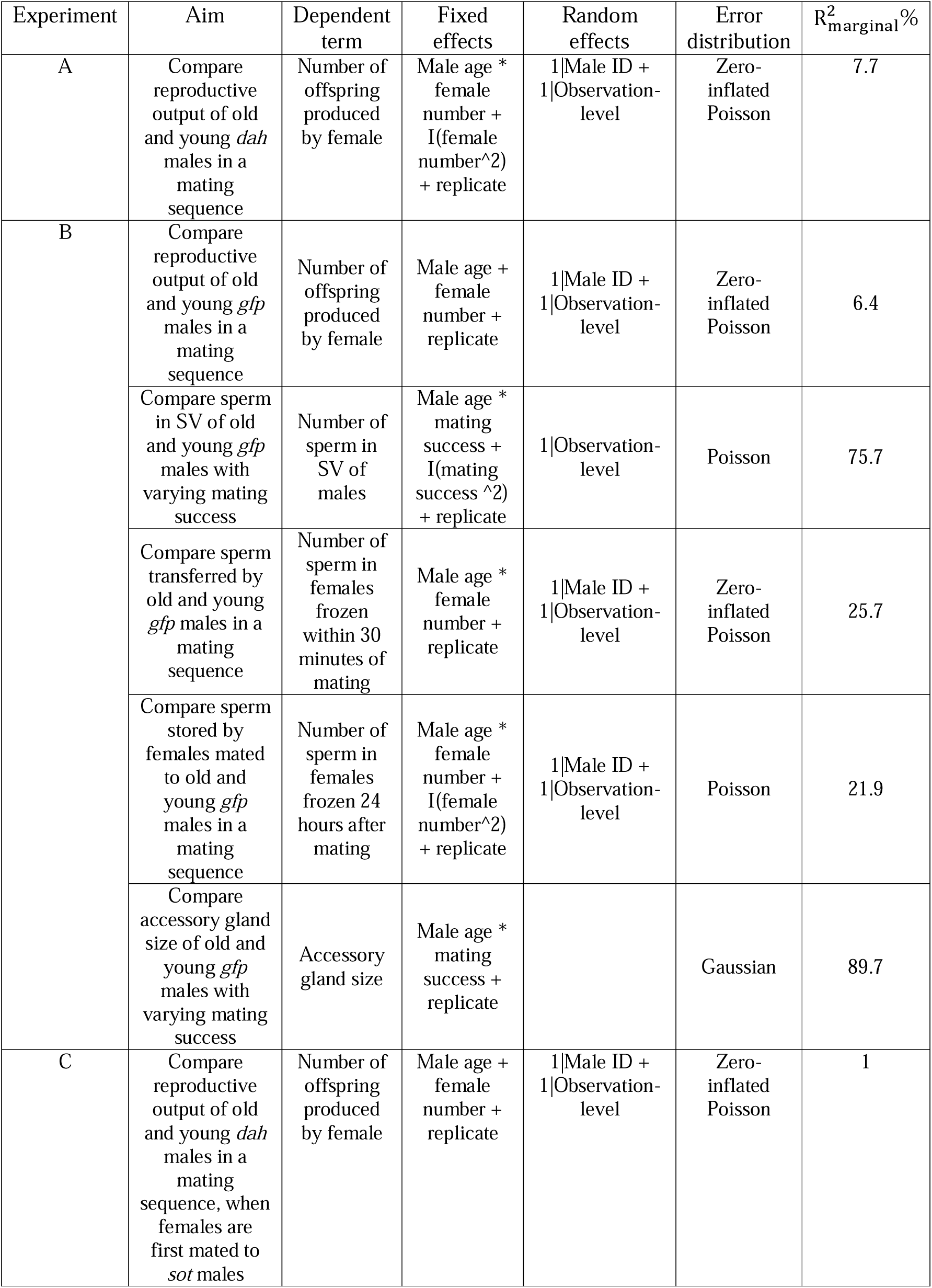
Descriptions of the best-fit models (final) models for each analysis in our study, with details on model aims, dependent and fixed terms, random effects, model error structure, and marginal variance explained. Model outputs of each best-fit model can be found in Tables S3-S9 below.

## Appendix 1 Scenarios for ‘male age x female number’ interactions

Some possible scenarios (Appendix figure 1) for how male age could interact with female number in a male’s multiple-mating sequence, to affect male reproductive output. We assume that male reproductive output (*W*) is a negative function of progression through the mating sequence, such that *W*(*i*) < *W*(*i*+1), where *i* is the *i*th female in a mating sequence. For simplicity we assume that the function is linear; i.e. *W* = −*ax* + *b*, where x is the order of female in the mating sequence, a is the slope, and b is the intercept. The <u>null hypothesis</u> is that old and young males have the same size and quality of ejaculates, and allocate ejaculates in similar ways to females in their mating sequence, thus produce similar numbers of offspring across a mating sequence:

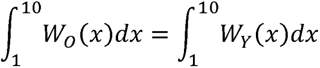

However, under scenarios of reproductive senescence (<u>scenario A</u>), the general prediction is that old males have overall lower total reproductive success than young males (i.e. lower intercepts; *b_O_<b_Y_*)).However, old and young males do not differ in their slopes (*a_O_ =* a_Y_) of decline through a mating sequence and allocate similar proportions of their ejaculate to each female. Here:

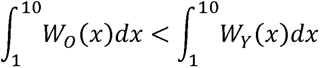

where *W_O_*and *W^Y^* are the reproductive output functions of old and young males over a mating sequence of 10 available females, respectively.

<u>Scenario B:</u> old males due to fewer mating opportunities in the future, might terminally invest in reproduction, allocating a higher proportion of ejaculate to early than late females compared to young males (i.e. |*a_O_*| > |*a_Y_* |). This would lead to relatively higher reproductive output of old males early in a mating sequence, but steeper slopes of decline through the mating sequence, compared to young males. <u>Scenario C:</u> Old males, due to having lower quantities of ejaculates, might more prudently allocate ejaculates to each female (i.e.|*a_O_* | < |*a_Y_*). This would lead old males to have lower intercepts but shallower slopes of decline in reproductive output than young males. <u>Scenario D:</u> Old males have smaller ejaculates than young males, however, they would transfer the same size of ejaculates to females as young males until old males run out of ejaculates. The point where old males run out of ejaculates would be sooner than that of young males, and old males would not have any reproductive output beyond this point.

**Appendix figure 1:**
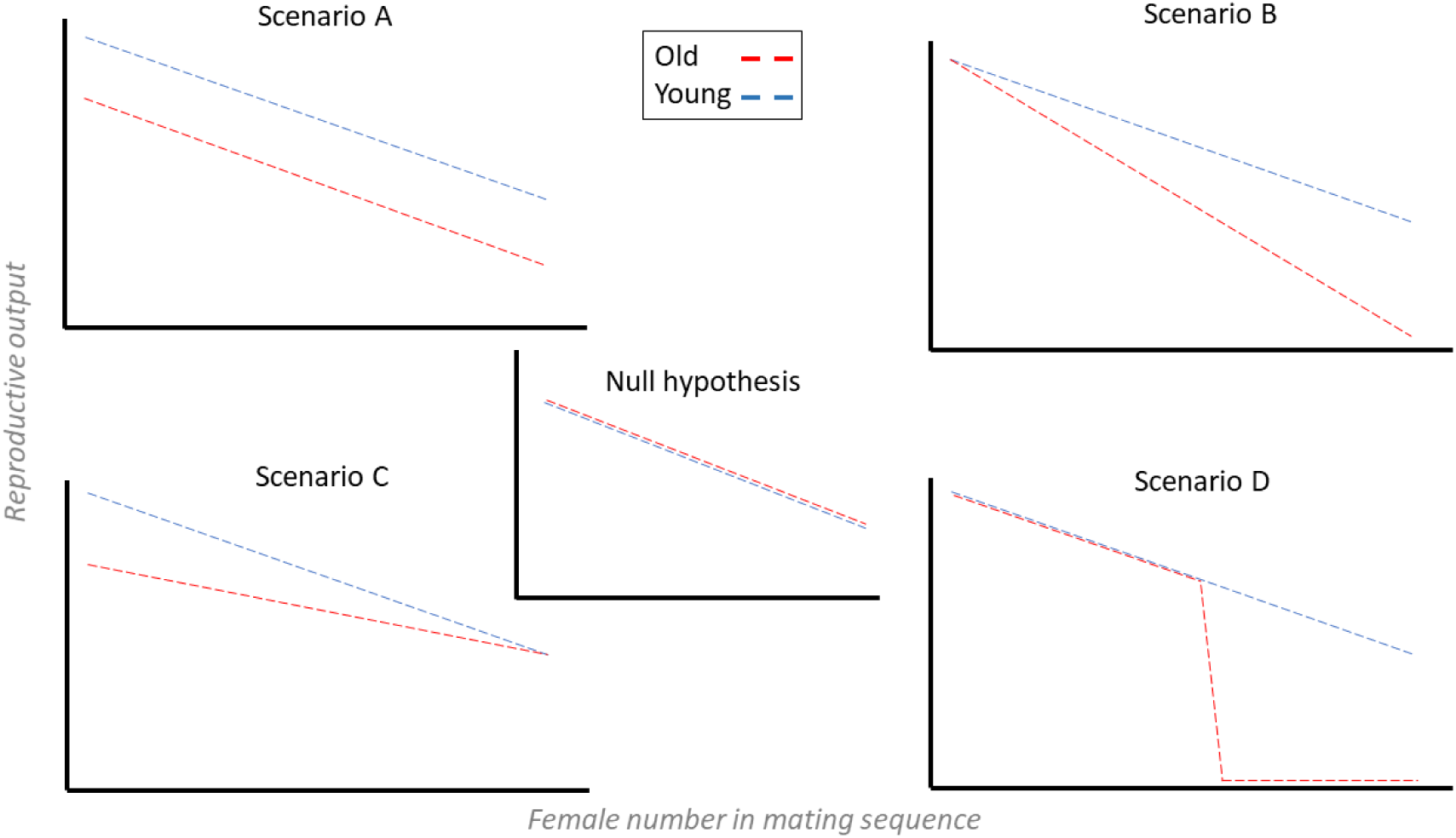
Possible scenarios for how advancing male age can interact with female order in the male’s mating sequence, to influence male reproductive output.

## Appendix 2 male mating success and latency

We compared the mating success (i.e. total number of females a male mated with) of young and old males in our three experiments. For this, we created a generalised linear model with Poisson error distribution, and included male mating success as our dependent variable. We modelled male age, experiment (A-C), and their two-way interaction as fixed effects, and observation-level ID as a random effect to account for overdispersion. We also compared male mating latency (time elapsed between male being paired with a female, and the start of copulation). For this, we modelled male mating latency as our dependent variable using an LMM with Gaussian error distribution. Male age, a linear and quadratic term of female number, their two-way interactions with male age, and experiment number, were included as fixed effects, with male ID as a random effect.

Old males consistently mated with fewer females compared to young males (z = 6.423, P < 0.001, Figure S4A). However, this difference was greater in Experiments B (age*experiment C: z = 3.491, P < 0.001) and C (age*experiment C: z = 2.027, P < 0.043) than Experiment A. Old males had a lower mating latency than young males early in the mating sequence but not later (age*female number: t = 3.159, P = 0.002, Figure S6B, S6C).

Ejaculate limitation seems to be an unlikely explanation for the lower mating success of old males, because they had more sperm in their SV and larger AGs than young males. Instead, the lower mating success of old males might be a consequence of their longer mating latencies. These longer latencies could be due to old males being less attractive to females or worse at courtship, thus females taking longer to accept mating with an old than a young male (e.g. Amin et al, 2012; Rezaei et al, 2015).

Furthermore, differences between old and young males in their mating latencies were greater in Experiment C (when females were first mated to *sot* males) than Experiment B (Figure S6C). This result suggests that females show stronger choice for young males than old males, when females have previously been mated compared to when females are virgins. Comparing male mating success and latency were not a part of our main aims; therefore, we have included these analyses in the appendix only.

## Appendix 3 Bateman’s gradients

We calculated Bateman’s gradients, i.e. the slope of the linear relationship between male mating success and reproductive success (Anthes et al, 2017), for old and young males used in Experiments A and C. This calculation was done to explore whether the opportunity for sexual selection is affected by intra-specific differences between males, namely age and seminal fluid availability. We first calculated the total mating success (sum of successful copulations by a male in his mating sequence), and total reproductive success (sum of offspring produced by all females that copulated with a male), for each male that mated in Experiments A and C. We then calculated the slopes of the linear regression between male mating success and reproductive success (Appendix figure 2). Visual inspection indicated that males in Experiment C had steeper slopes than males in Experiment A, suggesting that the opportunity for sexual selection might be greater when males are not seminal fluid limited. Additionally, young males had steeper gradients, thus a greater opportunity for pre-copulatory sexual selection, than old males. This result on Bateman’s gradients is only presented as an exploratory analysis to generate new hypotheses regarding how intra-specific variation between males might affect sexual selection. These analyses and results as such are not a part of our study aims.

**Appendix figure 2:**
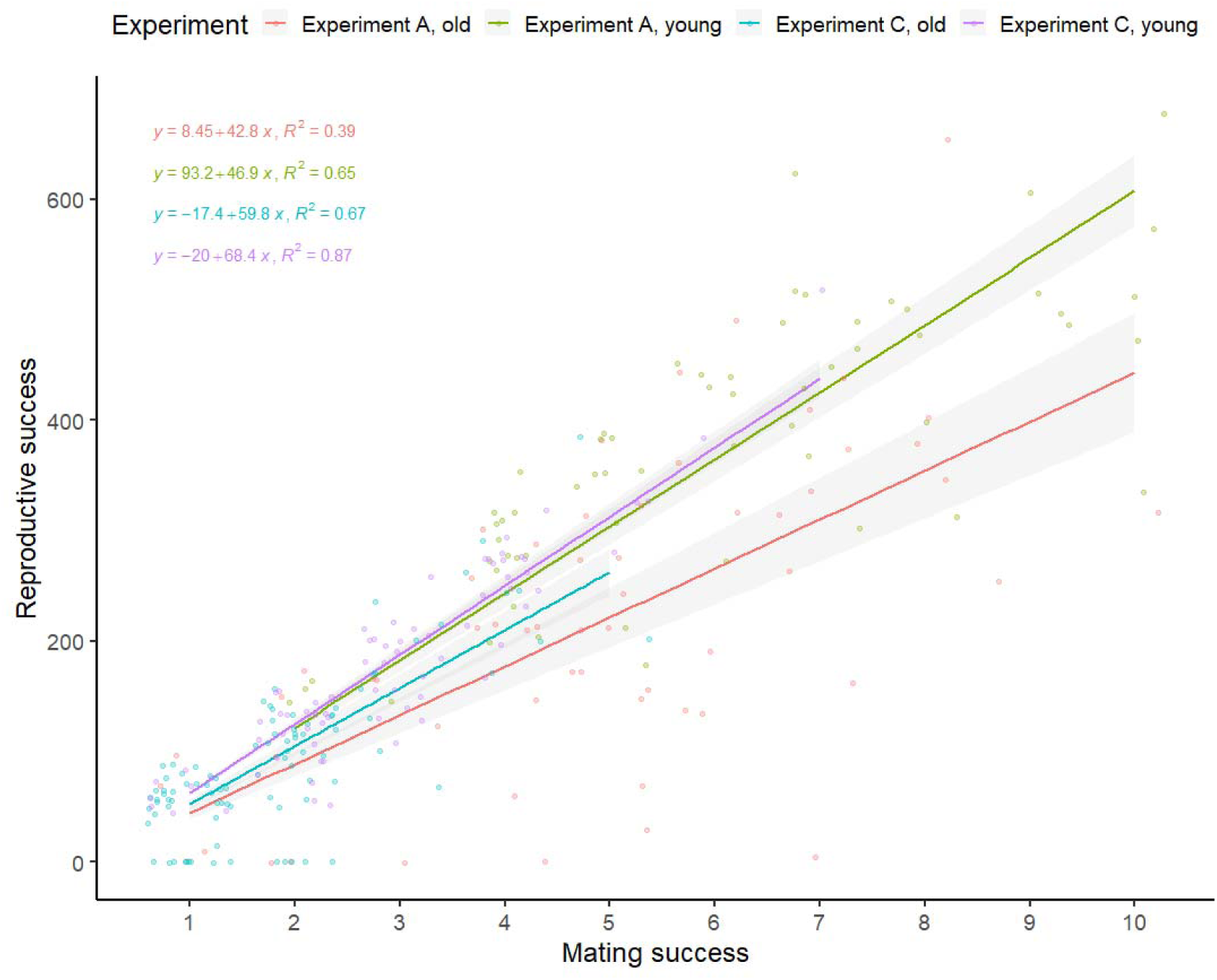
Bateman’s gradients showing the linear relationship between male mating success and reproductive success, for old and young males used in Experiments A and B. Dark lines show means, shaded regions show 95% C.I. Slopes and R^2^ values shown. Intercepts set to zero to allow direct comparisons of slopes. Each dot represents one male.

## Appendix 4 *son-of-tudor* crossing scheme

*Sot* males used in experiment C were generated by mating brown-eyed straight-winged homozygous *Tudor* females (backcrossed into *dah* background), to *dah* males. To ensure that this crossing scheme produced sterile flies, we used a sub-set of 120 non-experimental virgin *sot* males (3-4 days old), and kept them in bottles containing virgin *dah* females (∼20-30 males and females per bottle), for 14 days. None of the bottles had any larvae developing after 14 days, indicating that the males were sterile. We additionally dissected three randomly chosen *sot* males and examined their testes and AGs under a light microscope (Appendix figure 3), revealing the testes and SV to be shrunk and without sperm, but AGs appearing normal, indicating that the males were sterile but producing SF.

**Appendix figure 3:**
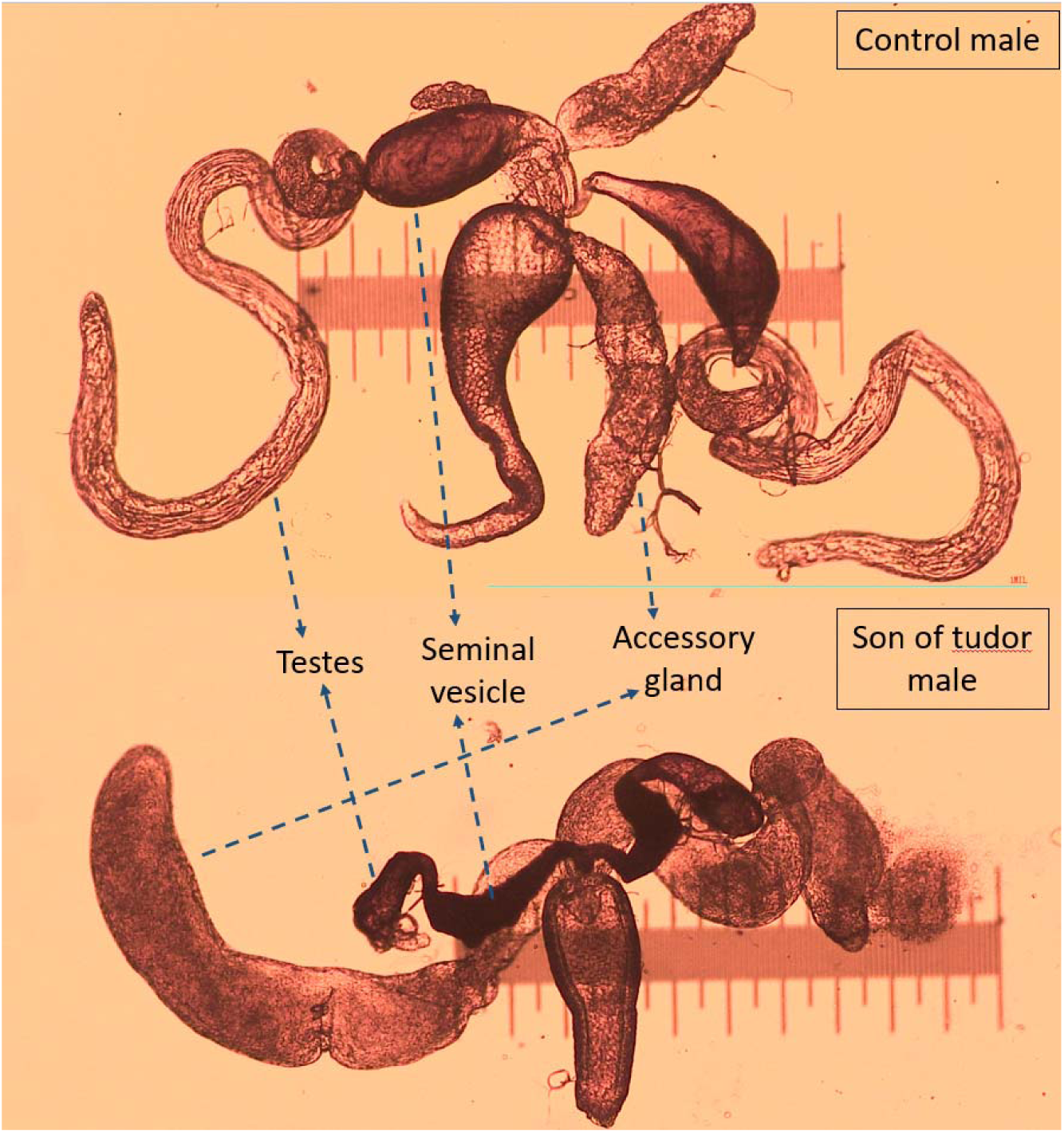
Disrupted testis and seminal vesicles (SV) with no sperm, but normal accessory glands (AG), in *sot* male, compared to control male. Scale = 1mm. Magnification = 40x (4x objective, 10x eyepiece).

## Appendix 5 male dissections and imaging

Males in experiment B were dissected, and later imaged to estimate sperm numbers stored in male seminal vesicles, and the size of male AGs. To dissect each male, three droplets of 50uL of PBS were placed on a slide covered with an aqueous solution of gelatine and chromium potassium sulphate. The reproductive tract of males (testes, accessory glands, seminal vesicles, and ejaculatory bulb) were separated from the rest of the body, by carefully pulling away the upper abdomen of the male using Inox Biology forceps. The male’s reproductive tract was washed in a second droplet of PBS and surrounding tissue separated. A clean sample of only reproductive tissue was placed on the third droplet. The accessory glands were then separated from the rest of the reproductive tract using microneedles (0.1mm thick), and placed on a new slide which had a measuring scale, inside a droplet of 5uL of PBS. The two accessory glands of each male were immediately imaged without a coverslip on, using the brightfield setting on a Nikon Eclipse 50i microscope with magnification of 4x (objective) and 10x (eyepiece), with each image calibrated to the scale of 1mm. Then, the male’s seminal vesicles were punctured, and sperm present in both seminal vesicles were carefully spread in the droplet of PBS using microneedles. A coverslip was then placed on this sample and then glued to the slide using rubber cement. The sperm were imaged the following day using a 5x air objective on a Zeiss LSM880 confocal laser scanning unit microscope (laser strength = 20; pinhole size = 20.1; Laser wavelength = 488; Pixel strength = 1196 x 1196; objective magnification = 5x, eyepiece magnification = 10x) and Zen black software.

To measure the area of accessory glands on FIJI/ImageJ win32, we used the freehand selection tool to outline the area of each accessory gland separately, and then used the “measure” option under “analyze” to measure the area of each accessory gland. To estimate the number of sperm in the seminal vesicles of males, we used the find maxima plugin (Prominence = 5000, strict setting, output type = single point). The prominence was chosen as 5000 because this point was where the power function relationship between prominence and estimates of sperm number by the find maxima plugin, became linear and flat (Appendix figure 4A, based on visual inspection of 5 samples). To ensure robustness of sperm numbers in male SV estimated by find maxima, we manually counted (using cell counter on FIJI) sperm in eight images of dissected male SV. Repeatability between manual counts and find maxima estimates of sperm numbers in male SV was high (R^2^ = 0.99, Appendix figure 4B).

**Appendix figure 4:**
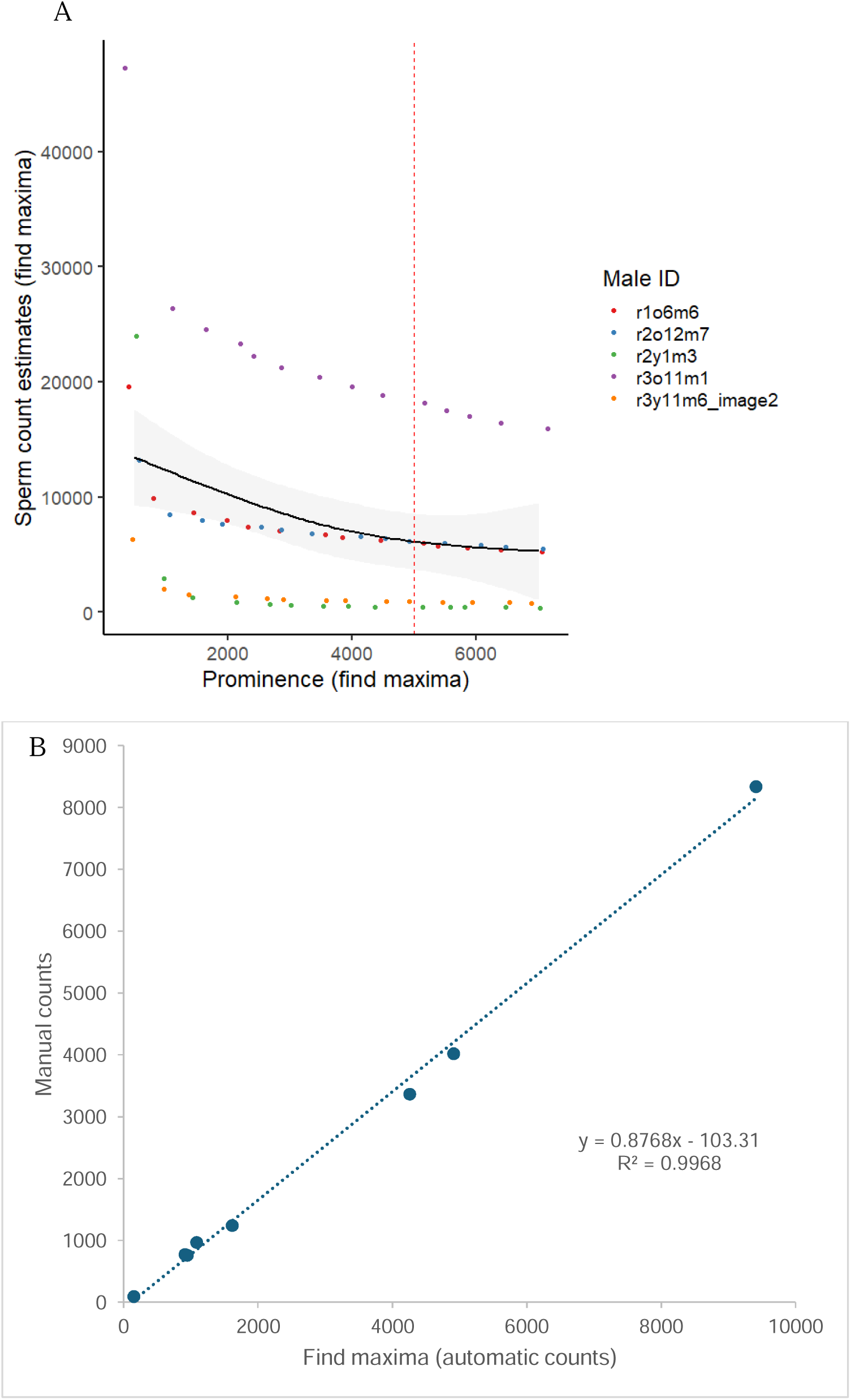
A. prominence on find maxima chosen as 5000 when estimating sperm stored in males, because this is where the power function relationship between prominence and estimates of sperm number by the find maxima plugin, became linear and flat based on visual inspection. B: High repeatability score of R^2^ = 0.99 between sperm numbers estimated manually (cell counter plugin) versus using find maxima plugin on FIJI.

## Appendix 6 Female dissections

Odd-numbered females (frozen after 24 hours) in a *gfp* male’s mating sequence, were dissected to count the number of sperm stored by a female in her long-term sperm storage organs. Even-numbered females in a *gfp* male’s mating sequence (frozen after 30 minutes), were dissected to count the number of sperm transferred by a male to a female.

To dissect frozen females, three droplets of 50uL of Phosphate buffer solution each, were placed on a slide coated with an aqueous solution of gelatine and chromium potassium sulphate. Each female to be dissected was placed on the first droplet, and her reproductive tract was removed gently by separating the last two abdominal segments from the rest of her body, using Inox Biology forceps.

Her reproductive organs (bursa, seminal receptacle, and spermathecae) were washed in the second droplet of PBS, and extra surrounding tissue removed. The reproductive organs were then placed on a third droplet. Here, using microneedles (0.1mm thick), the two spermathecae and seminal receptacle were gently spread and the sample covered with a coverslip. Rubber cement (fixogum) was used to glue the coverslip edges to the slide for imaging (Appendix 7) conducted on the following day.

## Appendix 7 sperm imaging and counts in females

For odd numbered females (frozen after 24 hours of mating), female reproductive tracts (spermathecae and seminal receptacle) fixed on a slide (described in Appendix 6) were imaged using a Nikon Eclipse50i fluorescence microscope (magnification = 10x objective, 10x eyepiece, wavelength = 480nm) with a chromix HD camera, under UV light from a CoolLED pe300 light source. The number of sperm heads (which appeared as fluorescent green under UV light) were later counted manually from images, using the cell counter plugin on FIJI/ImageJ version win32 (Schindelin et al, 2012). The GFP label in *gfp* flies is expressed at the *Mst35Ba* and *Mst35Bb* loci (Manier et al, 2010). To ensure repeatability of the manual counts, an independent analyst (TLC) counted 30 randomly chosen samples blind to counts made by the first analyst. Repeatability between the counts by the two analysts was high (R^2^ = 0.98, Appendix figure 5A). For even numbered females (frozen within 30 mins of mating), the bursa, spermathecae, and seminal receptacle, were imaged using a 5x air objective on a Zeiss LSM880 confocal laser scanning unit microscope (laser strength = 15; pinhole size = 34.1; laser wavelength = 488nm; pixel strength = 1196 x 1196, objective magnification = 5x, eyepiece magnification = 10x), and the Zen black v14.0.18.201 software. Images were analysed with ImageJ win32, and the find maxima plugin (with prominence = 10000, strict setting, output type = single point) was used to estimate the number of fluorescent green sperm heads in each sample. The prominence was chosen as 10000 because this was the point where the power function relationship between prominence and estimates of sperm number by the find maxima plugin, became linear and flat (Appendix figure 6, based on 8 samples, following Jahn et al (2021)). To ensure that sperm numbers estimated by the find maxima plugin were robust, we tested for repeatability between sperm number counted manually using the cell counter plugin, and sperm number estimated by the find maxima plugin. This repeatability test was done on a subset of 24 randomly chosen samples of even-numbered females (repeatability: R^2^ = 0.98, Appendix figure 5B). Examples of images of dissected individuals described in Appendix 5-7 are provided in Appendix figure 7.

**Appendix figure 5:**
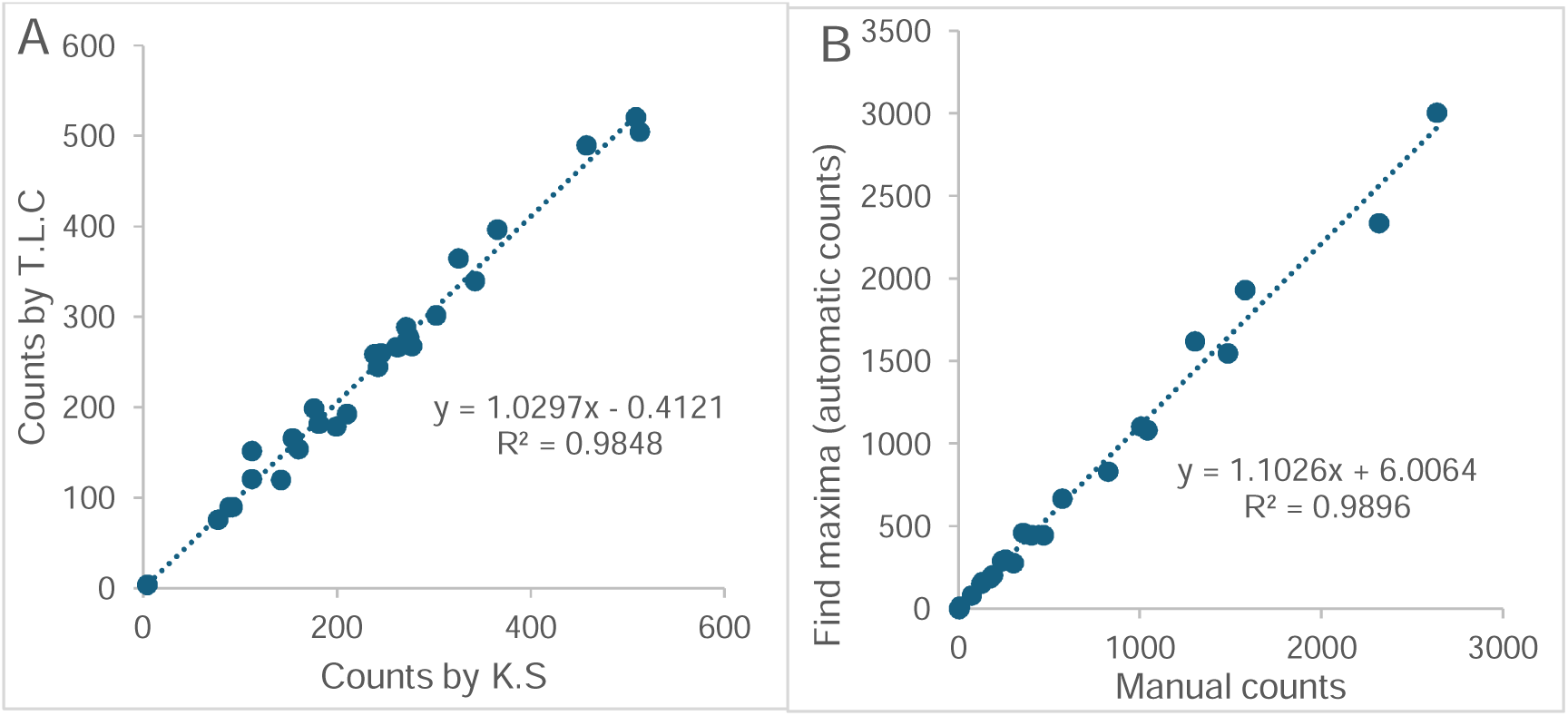
A: high repeatability score of R^2^ = 0.98 between two analysts, when counting sperm numbers stored in odd-numbered females after 24 hours, using the cell counter plugin. B: high repeatability score of R^2^ = 0.98, between sperm numbers when sperm are manually counted using the cell counter plugin, versus estimated using the find maxima plugin, for sperm transferred to even-numbered females.

**Appendix figure 6:**
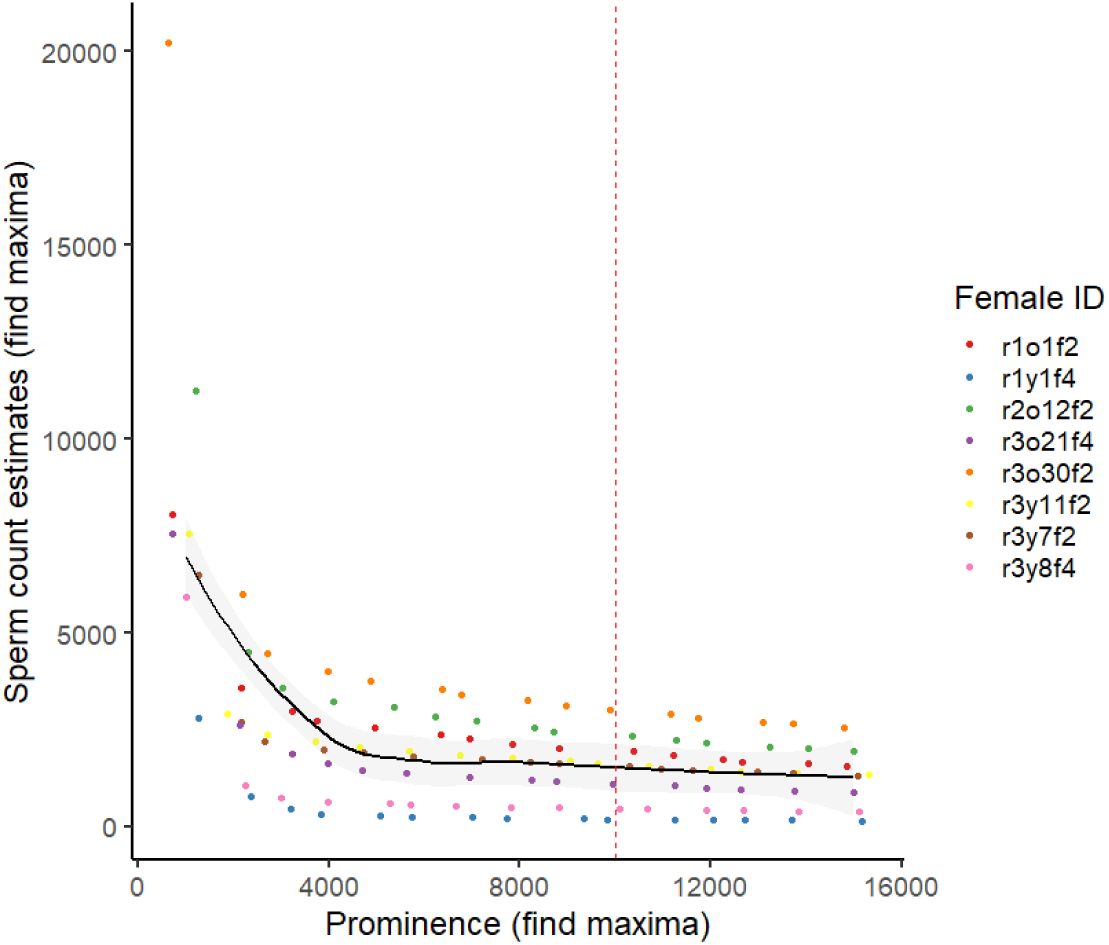
prominence on find maxima chosen as 10000, when estimating sperm stored in females, because this is where the power function relationship between prominence and estimates of sperm number by the find maxima plugin, became linear and flat based on visual inspection.

**Appendix figure 7:**
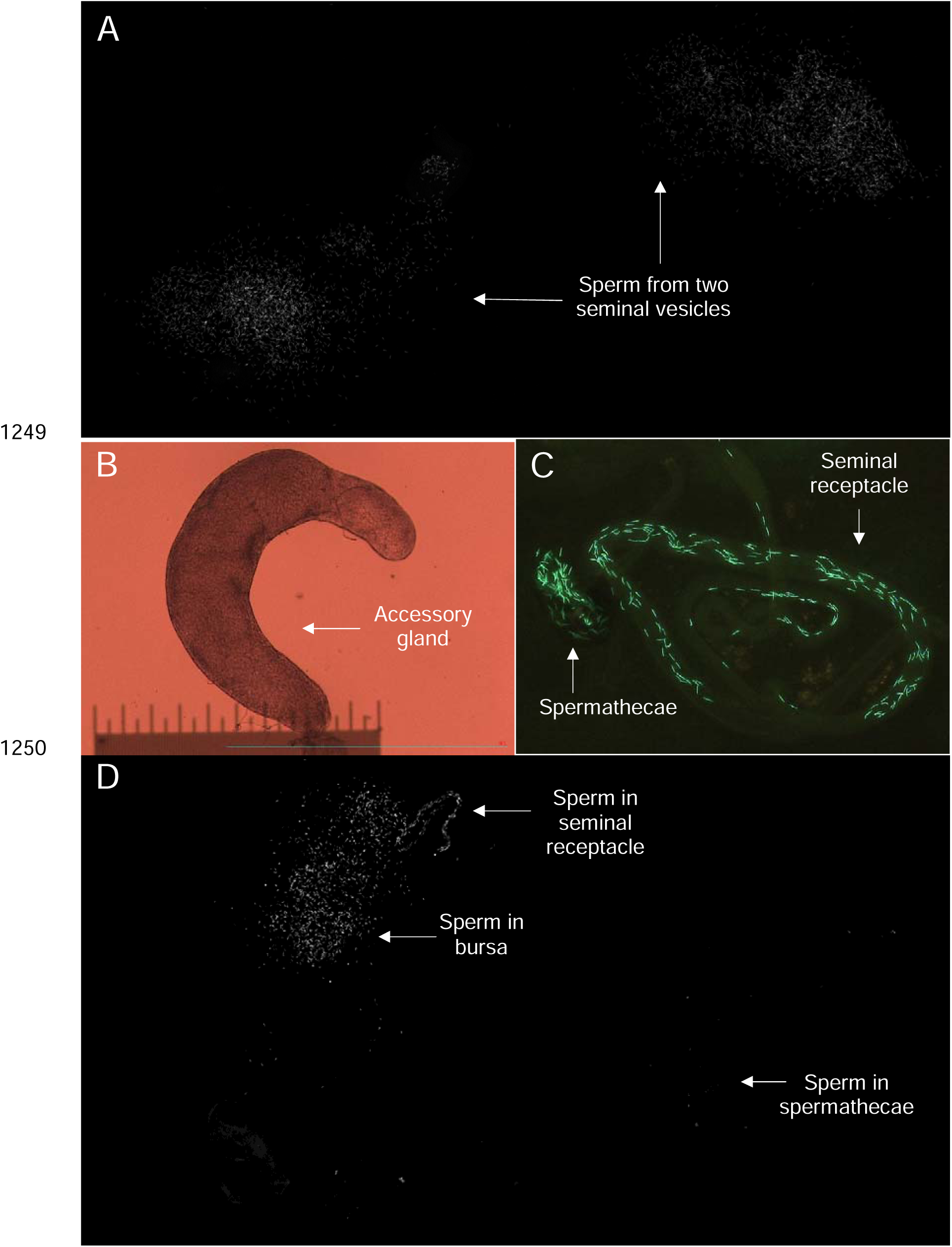
Examples of images from dissected individuals in Experiment B. **A.** sperm in seminal vesicles of males. **B.** accessory gland in male. **C.** sperm in long-term storage organs of even-numbered female after 24 hours. **D.** sperm transferred to odd-numbered female.

## Notes

https://osf.io/5z7m3/

### Competing Interest Statement

The authors have declared no competing interest.

