## Supplementary information for "Reproductive output of old polygynous males is limited by seminal fluid, not sperm number"

### Supplementary figures (S1-S7)

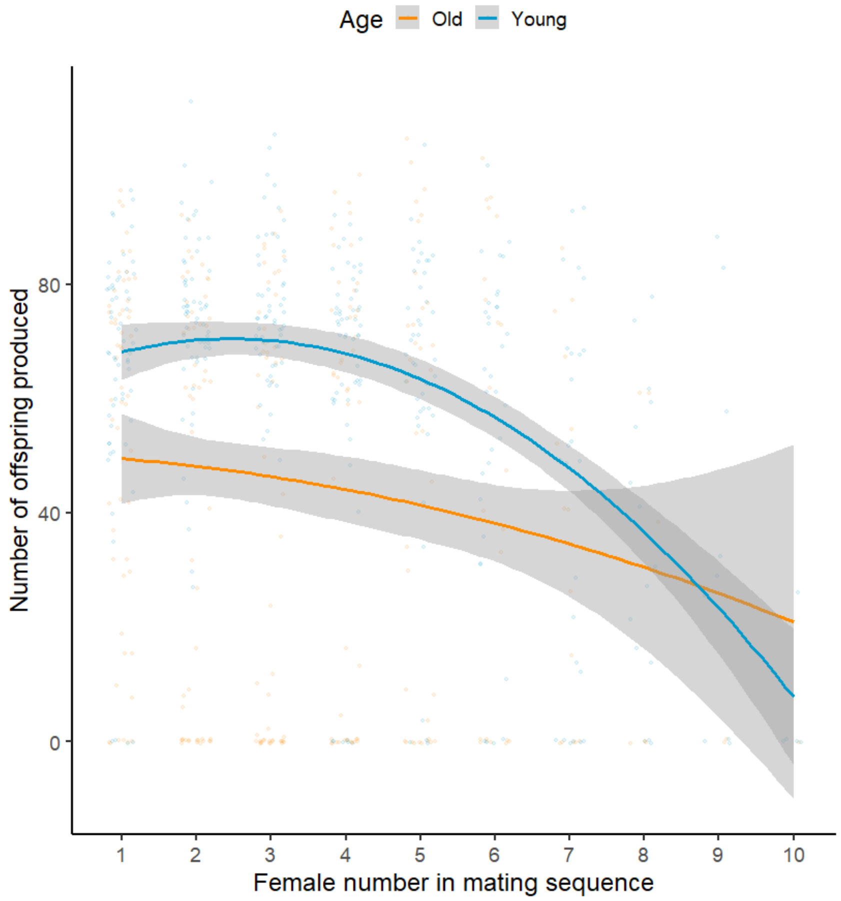


Fig S1: Significant interaction between male age and female number in a male's mating sequence, to affect the number of offspring produced by a female over 24 hours of egg laying, in Experiment A. Old males produce fewer offspring than young males only early on in a mating sequence. Means and 95% C.I. shown.

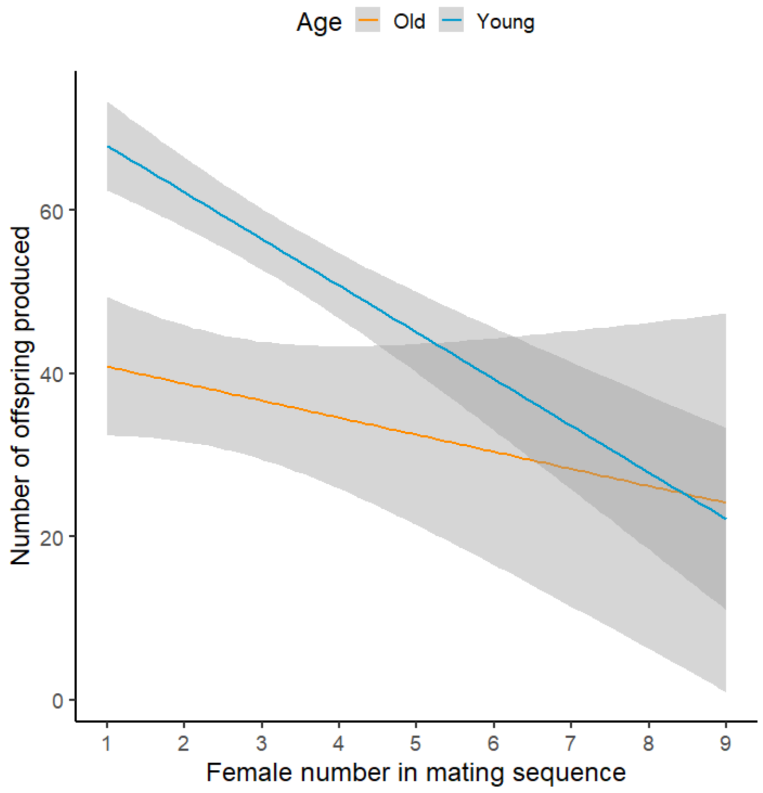


Fig S2: Significant effect of male age and female number in a male’s mating sequence, to affect the number of offspring produced by a female over 24 hours of egg laying, in Experiment B. Old males produce fewer offspring than young males, and males produce fewer offspring with females later than earlier in a mating sequence. Means and 95% C.I. shown.

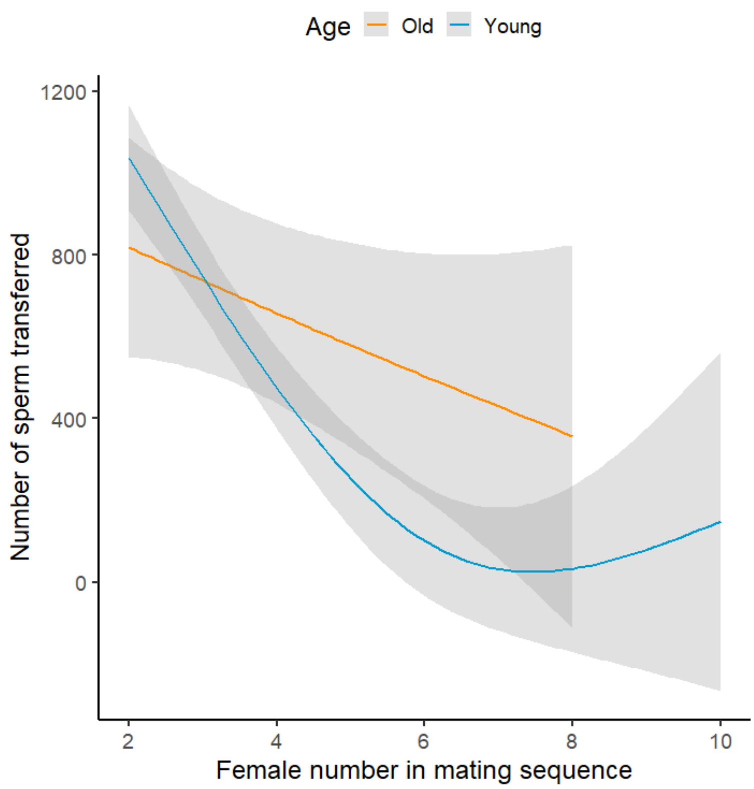


Figure S3: Effect of male age and female number in a male's mating sequence, on the number of sperm transferred by the male to the female, in experiment B. Means and 95% C.I. shown.

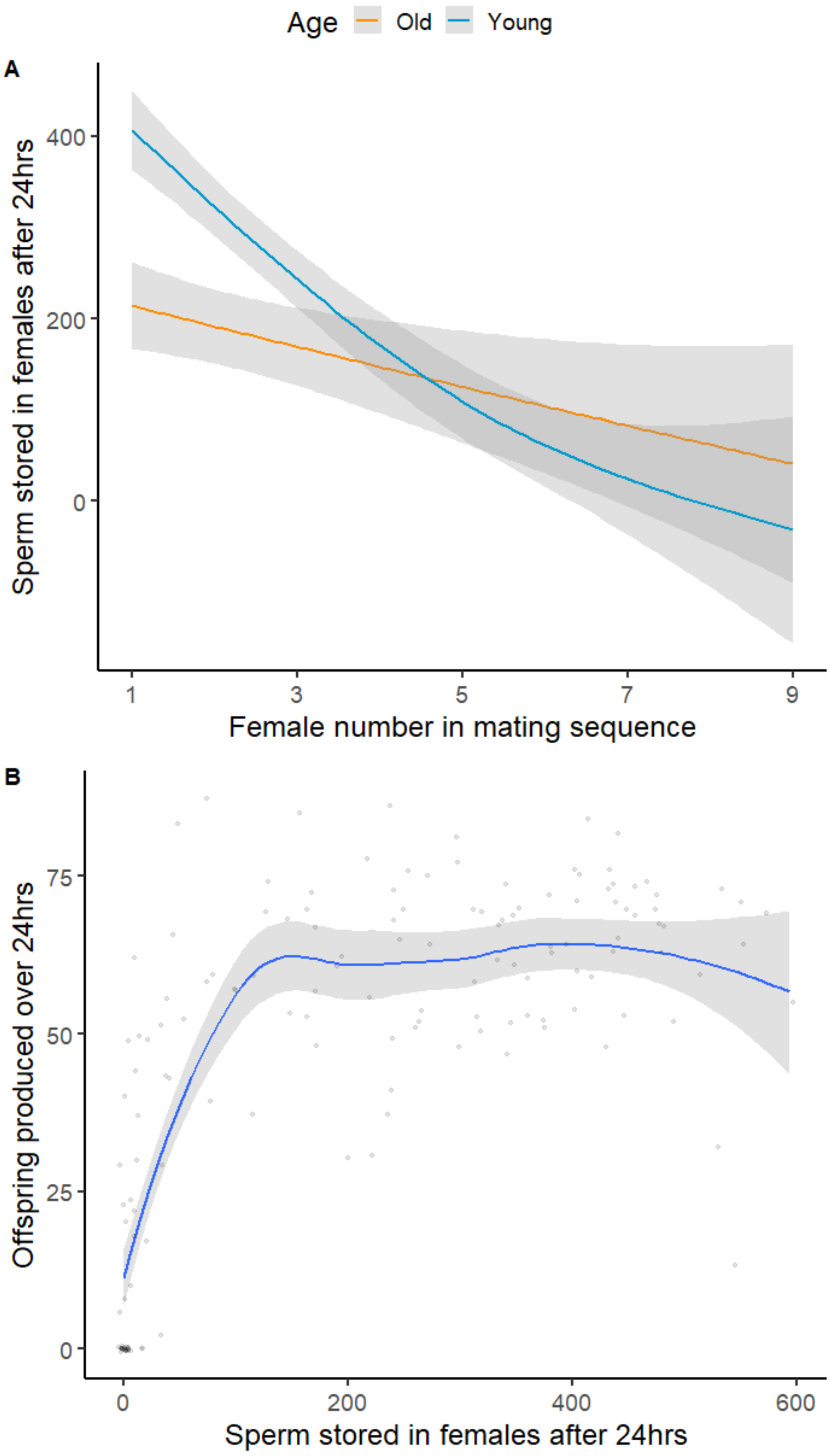


Figure S4: Experiment B. 4A- Effect of male age and female number in a male’s mating sequence, on the number of sperm stored by mated females after 24 hours of egg laying. 4B- Co-variance between number of sperm stored in odd-numbered females after 24 hours and the number of offspring produced by these females over 24 hours. Plot created using a loess smooth, to illustrate the non-linear (asymptotic) relationship. Means and 95% C.I. shown.

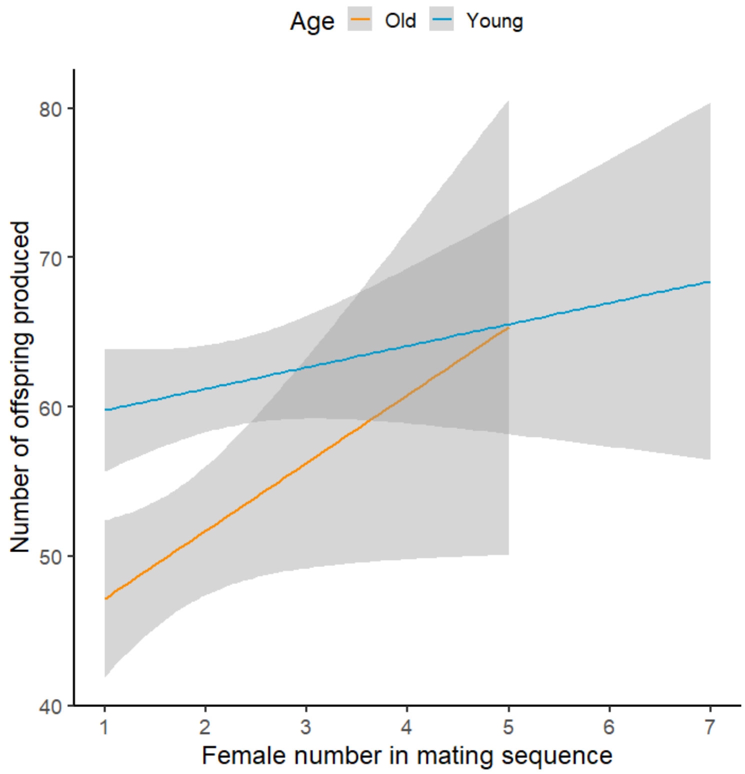


Fig S5: No significant effect of male age or female number in a male’s mating sequence, to affect the number of offspring produced by a female over 24 hours of egg laying, in Experiment C. Females, prior to focal mating with old or young *dah* males, were first mated with *sot* males to provide females with “extra” seminal fluid. Means and 95% C.I. shown.

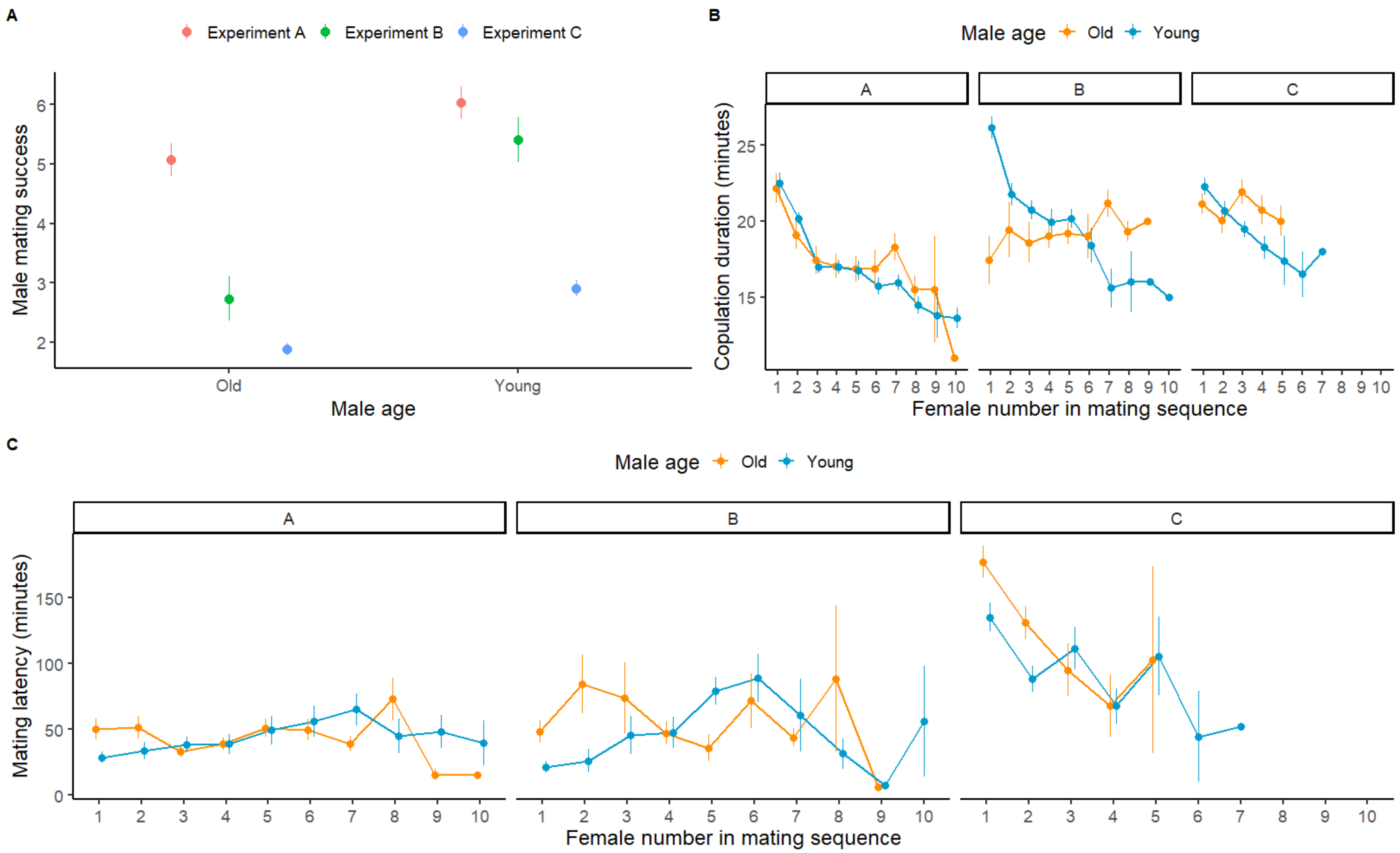


Figure S6: **A**. Mating success of focal old and young males used in our three experiments. **B.** Copulation duration of focal old and young males across Experiments A-C. **C.** Mating latency of focal young and old males across experiments A, B, and C. Means and SE shown.

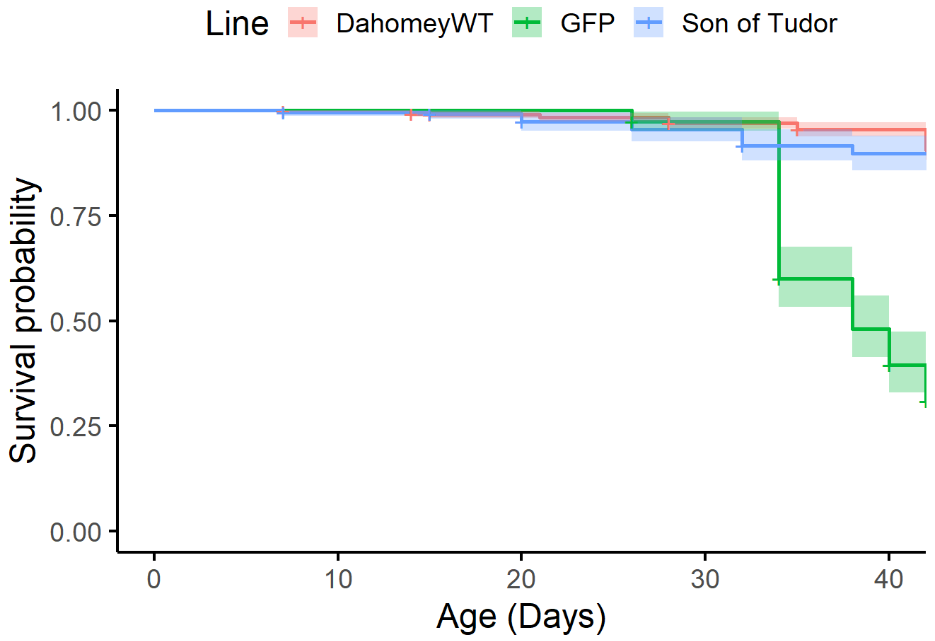


Figure S7: Age-dependent survival probability of males used in our study, from three distinct lines of *Drosophila melanogaster: dah, gfp, and sot*. *Gfp* males experienced higher mortality than *sot* or *dah* males. Means and 95% C.I. shown.

### Supplementary Tables (S1-S9)

Table S1: Effects of male age and female number in a male's mating sequence, on the number of offspring produced by mated females in Experiment A. Model constructed with zero inflated Poisson error distribution. Two-way interaction model used to interpret interaction only, main-effects model used to interpret main-effects only when two-way interaction is non-significant. Effects of interest highlighted in grey, significant P values of interest in bold.

Two-way interaction model

| Fixed effects | Estimate | SE | z | P |
|---|---|---|---|---|
| (Intercept) | 3.813 | 0.073 | 52.250 | <0.001 |
| I(Female number^2) | -0.014 | 0.004 | -3.430 | 0.001 |
| Age (Young) | 0.318 | 0.075 | 4.240 | <0.001 |
| Female number | 0.109 | 0.033 | 3.300 | 0.001 |
| RepA2 | 0.116 | 0.039 | 2.990 | 0.003 |
| Age (Young)*Female number | -0.043 | 0.019 | -2.240 | **0.025** |

| Random effects | Variance | SD |
|---|---|---|
| Male ID | 0.006 | 0.078 |
| Observation level | 0.146 | 0.382 |

Table S2: Effects of male age and female number in a male's mating sequence, on the number of offspring produced by mated females in Experiment B. Model constructed with zero inflated Poisson error distribution. Two-way interaction model used to interpret interaction only, main-effects model used to interpret main-effects only when two-way interaction is non-significant. Effects of interest highlighted in grey, significant P values of interest in bold.

Two-way interaction model

| Fixed effects | Estimate | SE | z | P |
|---|---|---|---|---|
| (Intercept) | 3.923 | 0.111 | 35.260 | <0.001 |
| Age (Young) | 0.266 | 0.126 | 2.110 | 0.035 |
| Female number | -0.037 | 0.030 | -1.240 | 0.216 |
| RepB2 | 0.112 | 0.108 | 1.040 | 0.299 |
| RepB3 | 0.082 | 0.092 | 0.890 | 0.376 |
| Age (Young)*Female number | -0.046 | 0.040 | -1.140 | 0.254 |

| Random effects | Variance | SD |
|---|---|---|
| Male ID | 0.000 | 0.000 |
| Observation level | 0.141 | 0.375 |

Main-effects model

| Fixed effects | Estimate | SE | z | P |
|---|---|---|---|---|
| (Intercept) | 3.979 | 0.101 | 39.540 | <0.001 |
| Age (Young) | 0.149 | 0.074 | 2.010 | **0.044** |
| Female number | -0.064 | 0.020 | -3.230 | **0.001** |
| RepCR2 | 0.124 | 0.108 | 1.150 | 0.250 |
| RepCR3 | 0.083 | 0.093 | 0.900 | 0.370 |

Table S3: Effects of male age and male mating success, on the number of sperm accumulated in seminal vesicles of males in experiment B (one data point per male). Model constructed with Poisson error distribution. Two-way interaction model used to interpret interaction only, main-effects model used to interpret main-effects only when two-way interaction is non-significant. Effects used for interpretation highlighted in grey, significant P values of interest in bold.

Two-way interaction model

| Fixed effects | Estimate | SE | z | P |
|---|---|---|---|---|
| (Intercept) | 9.632 | 0.165 | 58.350 | <0.001 |
| I(Mating success^2) | 0.035 | 0.007 | 4.990 | <0.001 |
| Age (Young) | -1.483 | 0.157 | -9.420 | <0.001 |
| Mating success | -0.377 | 0.062 | -6.120 | <0.001 |
| RepB2 | 0.144 | 0.158 | 0.910 | 0.363 |
| RepB3 | 0.514 | 0.143 | 3.600 | <0.001 |
| Age (Young)*Mating success | -0.166 | 0.036 | -4.630 | **<0.001** |

| Random effects | Variance | SD |
|---|---|---|
| Observation level | 0.179 | 0.423 |

Table S4: Effects of male age and female number in a male's mating sequence, on the number of sperm transferred by males to mated females in Experiment B. Model constructed with zero inflated Poisson error distribution. Two-way interaction model used to interpret interaction only, main-effects model used to interpret main-effects only when two-way interaction is non-significant. Effects used for interpretation highlighted in grey, significant P values of interest in bold.

Two-way interaction model

| Fixed effects | Estimate | SE | z | P |
|---|---|---|---|---|
| (Intercept) | 4.866 | 0.569 | 8.554 | <0.001 |
| Age (Young) | 2.433 | 0.594 | 4.099 | <0.001 |
| Female number | -0.033 | 0.115 | -0.288 | 0.773 |
| RepB2 | 0.182 | 0.526 | 0.346 | 0.730 |
| RepB3 | 0.675 | 0.461 | 1.464 | 0.143 |
| Age (Young)*Female number | -0.414 | 0.142 | -2.905 | **0.004** |

| Random effects | Variance | SD |
|---|---|---|
| Male ID | 0.631 | 0.795 |
| Observation level | 1.421 | 1.192 |

Table S5: Effects of male age and female number in a male's mating sequence, on the number of sperm transferred stored by mated females in Experiment B, after 24 hours of egg laying. Model constructed with Poisson error distribution. Two-way interaction model used to interpret interaction only, main-effects model used to interpret main-effects only when two-way interaction is non-significant. Effects used for interpretation highlighted in grey, significant P values of interest in bold.

Two-way interaction model

| Fixed effects | Estimate | SE | z | P |
|---|---|---|---|---|
| (Intercept) | 3.488 | 0.706 | 4.939 | <0.001 |
| I(Female number^2) | -0.032 | 0.039 | -0.821 | 0.412 |
| Age (Young) | 2.700 | 0.644 | 4.195 | <0.001 |
| Female number | -0.053 | 0.326 | -0.162 | 0.872 |
| RepB2 | -0.304 | 0.651 | -0.467 | 0.641 |
| RepB3 | 0.680 | 0.573 | 1.188 | 0.235 |
| Age (Young)*Female number | -0.373 | 0.184 | -2.030 | **0.042** |

| Random effects | Variance | SD |
|---|---|---|
| Male ID | 0.746 | 0.864 |
| Observation level | 4.244 | 2.060 |

Table S6: Effects of male age and male mating success, on the area of male accessory glands ($cm^2$) in experiment B (one data point per male). Model constructed with Gaussian error distribution. Two-way interaction model used to interpret interaction only. Effects used for interpretation highlighted in grey, significant P of interest values in bold.

Two-way interaction model

| Fixed effects | Estimate | SE | z | P |
|---|---|---|---|---|
| (Intercept) | 0.372 | 0.015 | 24.724 | <0.001 |
| Age (Young) | -0.161 | 0.014 | -11.677 | <0.001 |
| Mating success | -0.040 | 0.003 | -11.831 | <0.001 |
| RepB2 | -0.011 | 0.014 | -0.785 | 0.437 |
| RepB3 | 0.027 | 0.014 | 1.910 | 0.064 |
| Age (Young)*Mating success | 0.027 | 0.004 | 6.577 | **<0.001** |

Table S7: Effects of male age and female number in a male's mating sequence, on the number of offspring produced by mated females in Experiment C. Model constructed with zero inflated Poisson error distribution. Two-way interaction model used to interpret interaction only, main-effects model used to interpret main-effects only when two-way interaction is non-significant. Effects of interest highlighted in grey, significant P values of interest in bold.

Two-way interaction model

| Fixed effects | Estimate | SE | z | P |
|---|---|---|---|---|
| (Intercept) | 3.953 | 0.059 | 66.540 | <0.001 |
| Age (Young) | 0.154 | 0.074 | 2.080 | 0.038 |
| Female number | 0.055 | 0.029 | 1.940 | 0.053 |
| RepC2 | 0.038 | 0.034 | 1.120 | 0.264 |
| Age (Young)*Female number | -0.050 | 0.034 | -1.460 | 0.145 |

| Random effects | Variance | SD |
|---|---|---|
| Male ID | 0.000 | 0.020 |
| Observation level | 0.083 | 0.288 |

Main-effects model

| Fixed effects | Estimate | SE | z | P |
|---|---|---|---|---|
| (Intercept) | 4.015 | 0.041 | 96.820 | <0.001 |
| Age (Young) | 0.059 | 0.036 | 1.650 | 0.099 |
| Female number | 0.020 | 0.016 | 1.300 | 0.193 |
| RepC2 | 0.037 | 0.034 | 1.080 | 0.280 |

Table S8a: sample sizes for number of successful copulations by old and young males with females in a mating sequence, for Experiments A-C. Table 1b: sample sizes for total mating success (i.e. sum of females a male mated with) of old and young males, in Experiments A-C.

| Table 1a | | | | |
|---|---|---|---|---|
| Male Age | Female number in mating sequence (i.e. female mating order) | Sample sizes of successful copulations | | |
| | | Experiment A | Experiment B | Experiment C |
| Old | Starting male sample size | 60 | 50 | 115 |
| Old | $1^{st}$ | 57 | 45 | 89 |
| Old | $2^{nd}$ | 52 | 21 | 54 |
| Old | $3^{rd}$ | 49 | 15 | 19 |
| Old | $4^{th}$ | 45 | 14 | 8 |
| Old | $5^{th}$ | 35 | 9 | 2 |
| Old | $6^{th}$ | 22 | 9 | |
| Old | $7^{th}$ | 15 | 6 | |
| Old | $8^{th}$ | 6 | 3 | |
| Old | $9^{th}$ | 2 | 1 | |
| Old | $10^{th}$ | 1 | | |
| Young | Starting male sample size | 60 | 28 | 85 |
| Young | $1^{st}$ | 60 | 27 | 74 |
| Young | $2^{nd}$ | 60 | 26 | 67 |
| Young | $3^{rd}$ | 57 | 26 | 45 |
| Young | $4^{th}$ | 54 | 22 | 23 |
| Young | $5^{th}$ | 42 | 19 | 5 |
| Young | $6^{th}$ | 33 | 15 | 2 |
| Young | $7^{th}$ | 25 | 5 | 1 |
| Young | $8^{th}$ | 15 | 2 | |
| Young | $9^{th}$ | 9 | 2 | |
| Young | $10^{th}$ | 5 | 2 | |

| Table 1b | | | | |
|---|---|---|---|---|
| Male Age | Male mating success | Sample sizes of males | | |
| | | Experiment A | Experiment B | Experiment C |
| Old | 0 | 3 | 5 | 26 |
| Old | 1 | 5 | 24 | 35 |
| Old | 2 | 3 | 6 | 35 |
| Old | 3 | 4 | 1 | 11 |
| Old | 4 | 10 | 5 | 6 |
| Old | 5 | 13 | 0 | 2 |
| Old | 6 | 7 | 3 | |
| Old | 7 | 9 | 3 | |
| Old | 8 | 4 | 2 | |
| Old | 9 | 1 | 1 | |
| Old | 10 | 1 | | |
| Young | 0 | 0 | 1 | 11 |
| Young | 1 | 0 | 1 | 7 |
| Young | 2 | 3 | 0 | 22 |
| Young | 3 | 3 | 4 | 22 |
| Young | 4 | 12 | 3 | 18 |
| Young | 5 | 9 | 4 | 3 |
| Young | 6 | 8 | 10 | 1 |
| Young | 7 | 10 | 3 | 1 |
| Young | 8 | 6 | 0 | 0 |
| Young | 9 | 4 | 0 | 0 |
| Young | 10 | 5 | 2 | 0 |

Table S9: Descriptions of the best-fit models (final) models for each analysis in our study, with details on model aims, dependent and fixed terms, random effects, model error structure, and marginal variance explained. Model outputs of each best-fit model can be found in Tables S3-S9 below.

| Experiment | Aim | Dependent term | Fixed effects | Random effects | Error distribution | $R^2_{marginal}$% |
|---|---|---|---|---|---|---|
| A | Compare reproductive output of old and young *dah* males in a mating sequence | Number of offspring produced by female | Male age * female number + I(female number^2) + replicate | 1\|Male ID + 1\|Observation-level | Zero-inflated Poisson | 7.7 |
| B | Compare reproductive output of old and young *gfp* males in a mating sequence | Number of offspring produced by female | Male age + female number + replicate | 1\|Male ID + 1\|Observation-level | Zero-inflated Poisson | 6.4 |
| | Compare sperm in SV of old and young *gfp* males with varying mating success | Number of sperm in SV of males | Male age * mating success + I(mating success ^2) + replicate | 1\|Observation-level | Poisson | 75.7 |
| | Compare sperm transferred by old and young *gfp* males in a mating sequence | Number of sperm in females frozen within 30 minutes of mating | Male age * female number + replicate | 1\|Male ID + 1\|Observation-level | Zero-inflated Poisson | 25.7 |
| | Compare sperm stored by females mated to old and young *gfp* males in a mating sequence | Number of sperm in females frozen 24 hours after mating | Male age * female number + I(female number^2) + replicate | 1\|Male ID + 1\|Observation-level | Poisson | 21.9 |
| | Compare accessory gland size of old and young *gfp* males with varying mating success | Accessory gland size | Male age * mating success + replicate | | Gaussian | 89.7 |
| C | Compare reproductive output of old and young *dah* males in a mating sequence, when females are first mated to *sot* males | Number of offspring produced by female | Male age + female number + replicate | 1\|Male ID + 1\|Observation-level | Zero-inflated Poisson | 1 |

### Appendix

#### Appendix 1: Scenarios for ‘male age x female number’ interactions

Some possible scenarios (Appendix figure 1) for how male age could interact with female number in a male’s multiple-mating sequence, to affect male reproductive output. We assume that male reproductive output (*W*) is a negative function of progression through the mating sequence, such that $W(i) < W(i+1)$ , where *i* is the *i*th female in a mating sequence. For simplicity we assume that the function is linear; i.e. $W = -ax + b$, where x is the order of female in the mating sequence, a is the slope, and b is the intercept. The <u>null hypothesis</u> is that old and young males have the same size and quality of ejaculates, and allocate ejaculates in similar ways to females in their mating sequence, thus produce similar numbers of offspring across a mating sequence:

$$\int_{1}^{10} W_O(x)dx = \int_{1}^{10} W_Y(x)dx$$

However, under scenarios of reproductive senescence (<u>scenario A</u>), the general prediction is that old males have overall lower total reproductive success than young males (i.e. lower intercepts; $b_O<b_Y$)). However, old and young males do not differ in their slopes ($a_O = a_Y$) of decline through a mating sequence and allocate similar proportions of their ejaculate to each female. Here:

$$\int_{1}^{10} W_O(x)dx < \int_{1}^{10} W_Y(x)dx$$

where $W_O$ and $W^Y$ are the reproductive output functions of old and young males over a mating sequence of 10 available females, respectively.

<u>Scenario B:</u> old males due to fewer mating opportunities in the future, might terminally invest in reproduction, allocating a higher proportion of ejaculate to early than late females compared to young males (i.e. $|a_O| > |a_Y|$). This would lead to relatively higher reproductive output of old males early in a mating sequence, but steeper slopes of decline through the mating sequence, compared to young males. <u>Scenario C:</u> Old males, due to having lower quantities of ejaculates, might more prudently allocate ejaculates to each female $(\text{i.e.}\, |a_O| < |a_Y)$. This would lead old males to have lower intercepts but shallower slopes of decline in reproductive output than young males. <u>Scenario D:</u> Old males have smaller ejaculates than young males, however, they would transfer the same size of ejaculates to females as young males until old males run out of ejaculates. The point where old males run out of ejaculates would be sooner than that of young males, and old males would not have any reproductive output beyond this point.

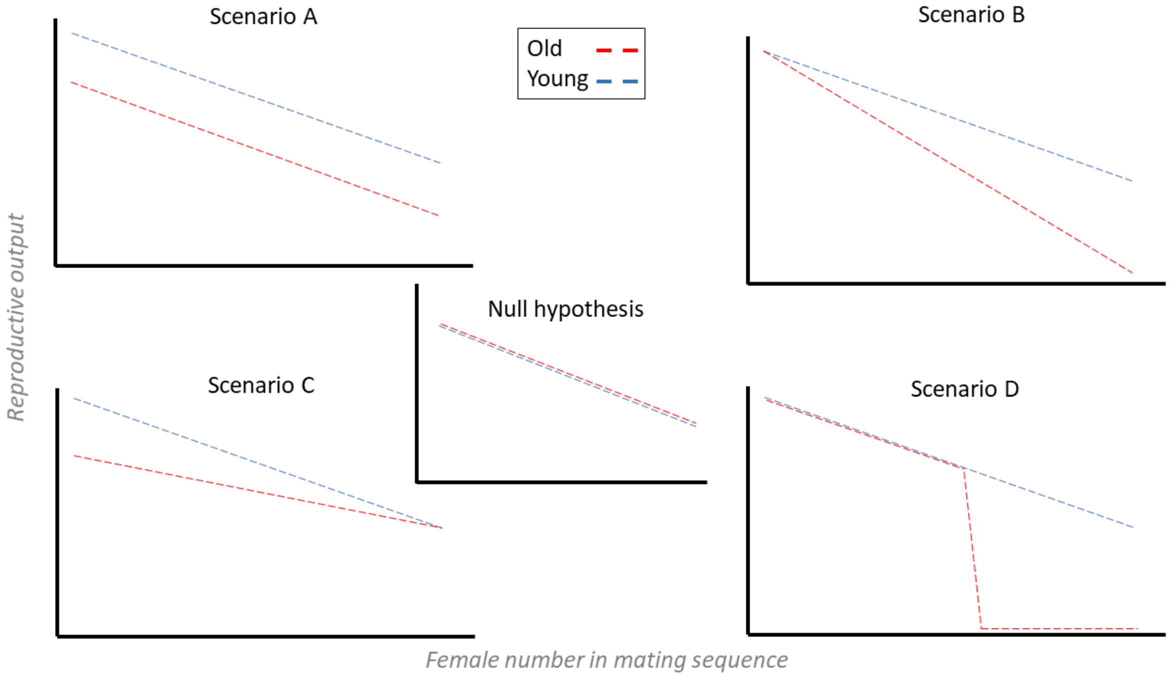


Appendix figure 1: Possible scenarios for how advancing male age can interact with female order in the male's mating sequence, to influence male reproductive output.

#### Appendix 2: male mating success and latency

#### Appendix 3: Bateman’s gradients

We calculated Bateman’s gradients, i.e. the slope of the linear relationship between male mating success and reproductive success (Anthes et al, 2017), for old and young males used in Experiments A and C. This calculation was done to explore whether the opportunity for sexual selection is affected by intra-specific differences between males, namely age and seminal fluid availability. We first calculated the total mating success (sum of successful copulations by a male in his mating sequence), and total reproductive success (sum of offspring produced by all females that copulated with a male), for each male that mated in Experiments A and C. We then calculated the slopes of the linear regression between male mating success and reproductive success (Appendix figure 2). Visual inspection indicated that males in Experiment C had steeper slopes than males in Experiment A, suggesting that the opportunity for sexual selection might be greater when males are not seminal fluid limited. Additionally, young males had steeper gradients, thus a greater opportunity for pre-copulatory sexual selection, than old males. This result on Bateman’s gradients is only presented as an exploratory analysis to generate new hypotheses regarding how intra-specific variation between males might affect sexual selection. These analyses and results as such are not a part of our study aims.

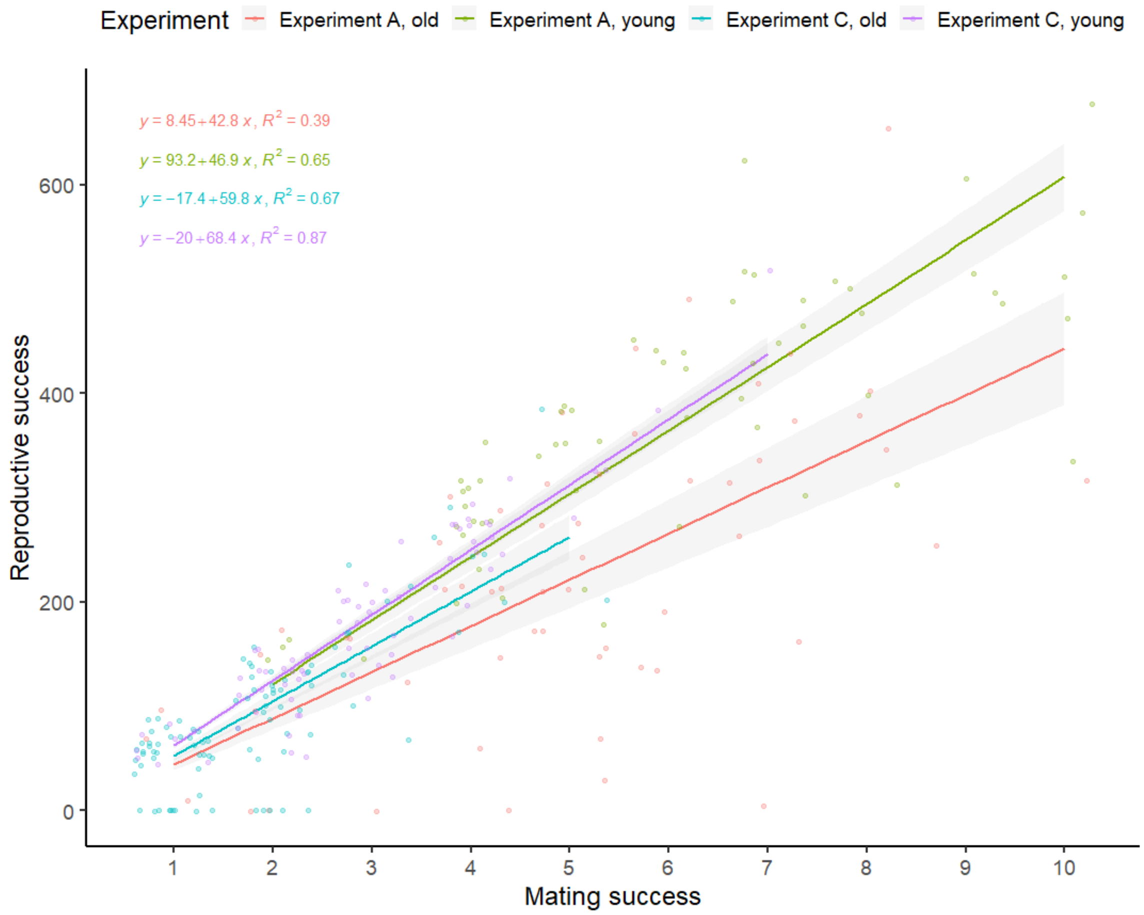


Appendix figure 2: Bateman’s gradients showing the linear relationship between male mating success and reproductive success, for old and young males used in Experiments A and B. Dark lines show means, shaded regions show 95% C.I. Slopes and $R^2$ values shown. Intercepts set to zero to allow direct comparisons of slopes. Each dot represents one male.

Appendix 4: *son-of-tudor* crossing scheme

*Sot* males used in experiment C were generated by mating brown-eyed straight-winged homozygous *Tudor* females (backcrossed into *dah* background), to *dah* males. To ensure that this crossing scheme produced sterile flies, we used a sub-set of 120 non-experimental virgin *sot* males (3-4 days old), and kept them in bottles containing virgin *dah* females (~20-30 males and females per bottle), for 14 days. None of the bottles had any larvae developing after 14 days, indicating that the males were sterile. We additionally dissected three randomly chosen *sot* males and examined their testes and AGs under a light microscope (Appendix figure 3), revealing the testes and SV to be shrunk and without sperm, but AGs appearing normal, indicating that the males were sterile but producing SF.

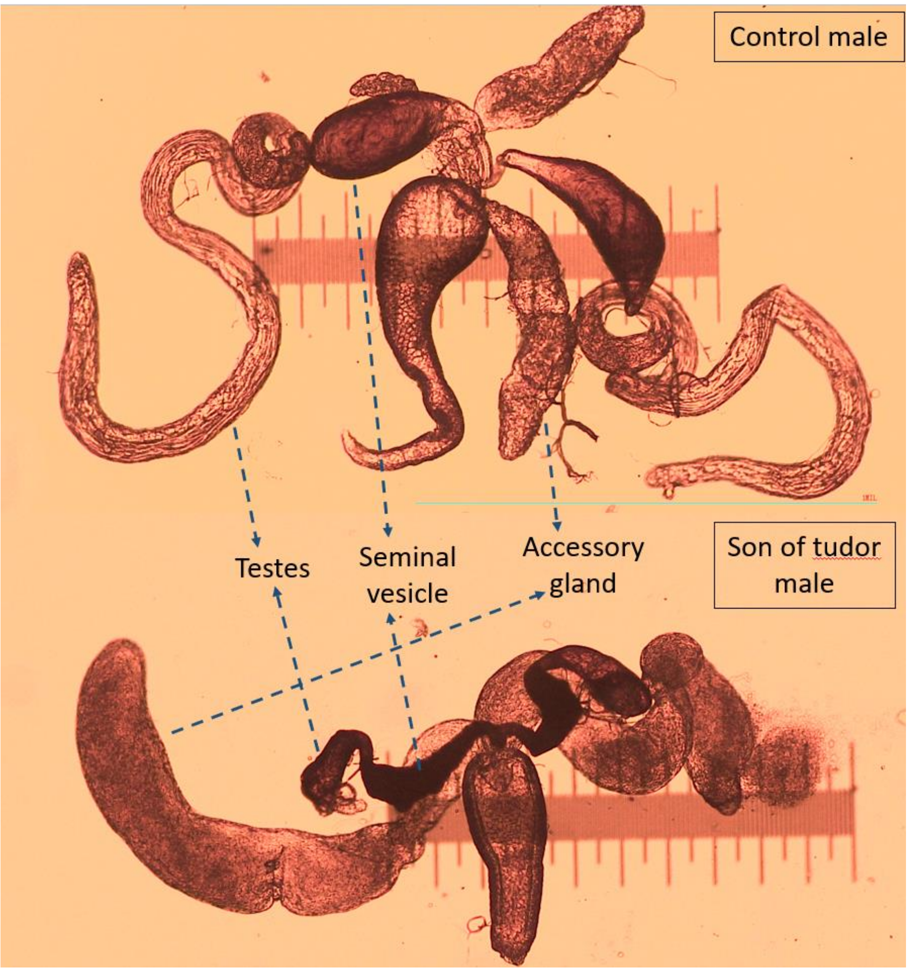


Appendix figure 3: Disrupted testis and seminal vesicles (SV) with no sperm, but normal accessory glands (AG), in *sot* male, compared to control male. Scale = 1mm. Magnification = 40x (4x objective, 10x eyepiece).

#### Appendix 5: male dissections and imaging

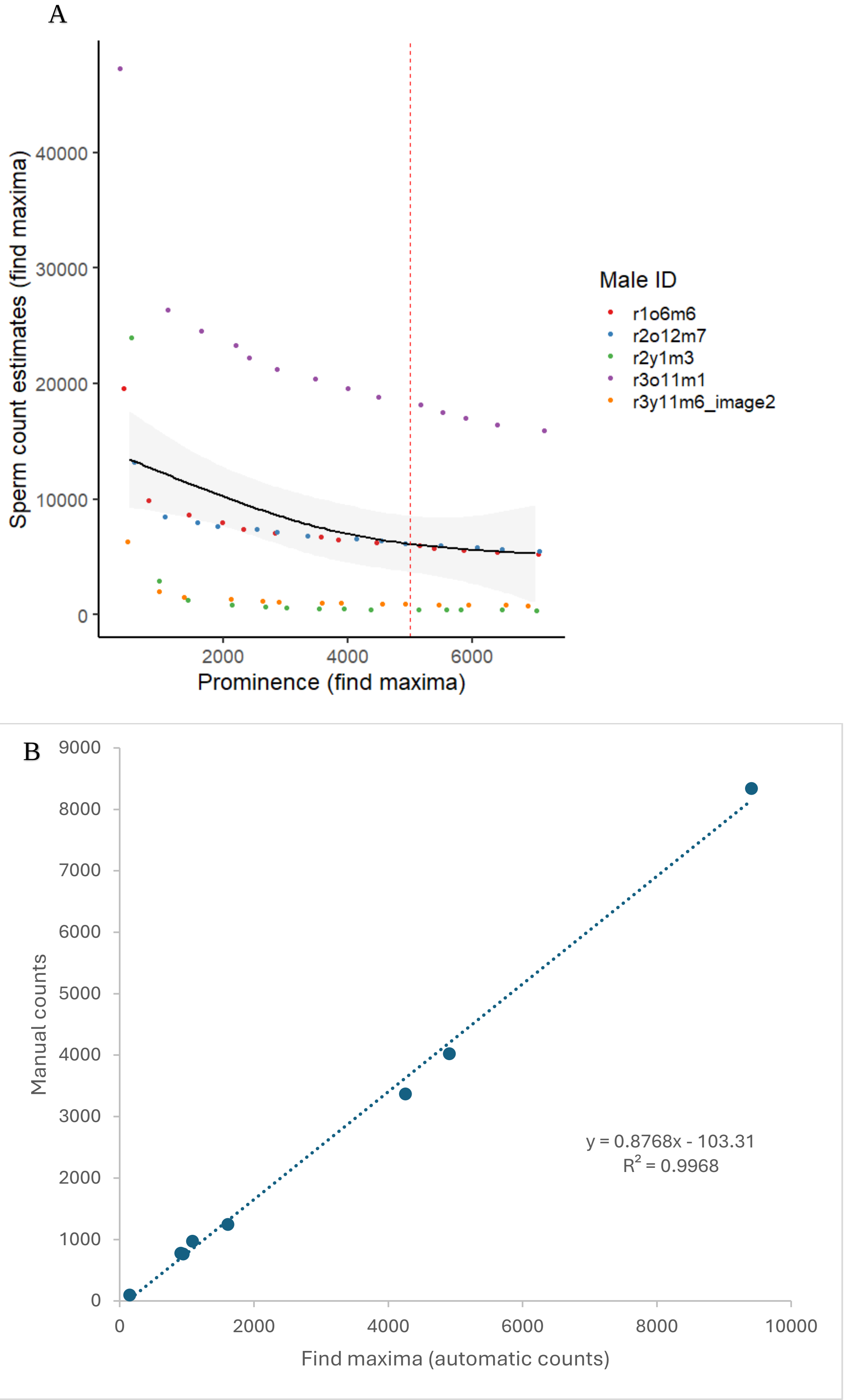


Appendix figure 4: A. prominence on find maxima chosen as 5000 when estimating sperm stored in males, because this is where the power function relationship between prominence and estimates of sperm number by the find maxima plugin, became linear and flat based on visual inspection. B: High repeatability score of $R^2 = 0.99$ between sperm numbers estimated manually (cell counter plugin) versus using find maxima plugin on FIJI.

#### Appendix 6: Female dissections

Odd-numbered females (frozen after 24 hours) in a *gfp* male's mating sequence, were dissected to count the number of sperm stored by a female in her long-term sperm storage organs. Even-numbered females in a *gfp* male's mating sequence (frozen after 30 minutes), were dissected to count the number of sperm transferred by a male to a female.

To dissect frozen females, three droplets of 50uL of Phosphate buffer solution each, were placed on a slide coated with an aqueous solution of gelatine and chromium potassium sulphate. Each female to be dissected was placed on the first droplet, and her reproductive tract was removed gently by separating the last two abdominal segments from the rest of her body, using Inox Biology forceps. Her reproductive organs (bursa, seminal receptacle, and spermathecae) were washed in the second droplet of PBS, and extra surrounding tissue removed. The reproductive organs were then placed on a third droplet. Here, using microneedles (0.1mm thick), the two spermathecae and seminal receptacle were gently spread and the sample covered with a coverslip. Rubber cement (fixogum) was used to glue the coverslip edges to the slide for imaging (Appendix 7) conducted on the following day.

#### Appendix 7: sperm imaging and counts in females

For odd numbered females (frozen after 24 hours of mating), female reproductive tracts (spermathecae and seminal receptacle) fixed on a slide (described in Appendix 6) were imaged using a Nikon Eclipse50i fluorescence microscope (magnification = 10x objective, 10x eyepiece, wavelength = 480nm) with a chromix HD camera, under UV light from a CoolLED pe300 light source. The number of sperm heads (which appeared as fluorescent green under UV light) were later counted manually from images, using the cell counter plugin on FIJI/ImageJ version win32 (Schindelin et al, 2012). The GFP label in *gfp* flies is expressed at the *Mst35Ba* and *Mst35Bb* loci (Manier et al, 2010). To ensure repeatability of the manual counts, an independent analyst (TLC) counted 30 randomly chosen samples blind to counts made by the first analyst. Repeatability between the counts by the two analysts was high ($R^2$ = 0.98, Appendix figure 5A). For even numbered females (frozen within 30 mins of mating), the bursa, spermathecae, and seminal receptacle, were imaged using a 5x air objective on a Zeiss LSM880 confocal laser scanning unit microscope (laser strength = 15; pinhole size = 34.1; laser wavelength = 488nm; pixel strength = 1196 x 1196, objective magnification = 5x, eyepiece magnification = 10x), and the Zen black v14.0.18.201 software. Images were analysed with ImageJ win32, and the find maxima plugin (with prominence = 10000, strict setting, output type = single point) was used to estimate the number of fluorescent green sperm heads in each sample. The prominence was chosen as 10000 because this was the point where the power function relationship between prominence and estimates of sperm number by the find maxima plugin, became linear and flat (Appendix figure 6, based on 8 samples, following Jahn et al (2021)). To ensure that sperm numbers estimated by the find maxima plugin were robust, we tested for repeatability between sperm number counted manually using the cell counter plugin, and sperm number estimated by the find maxima plugin. This repeatability test was done on a subset of 24 randomly chosen samples of even-numbered females (repeatability: $R^2$ = 0.98, Appendix figure 5B). Examples of images of dissected individuals described in Appendix 5-7 are provided in Appendix figure 7.

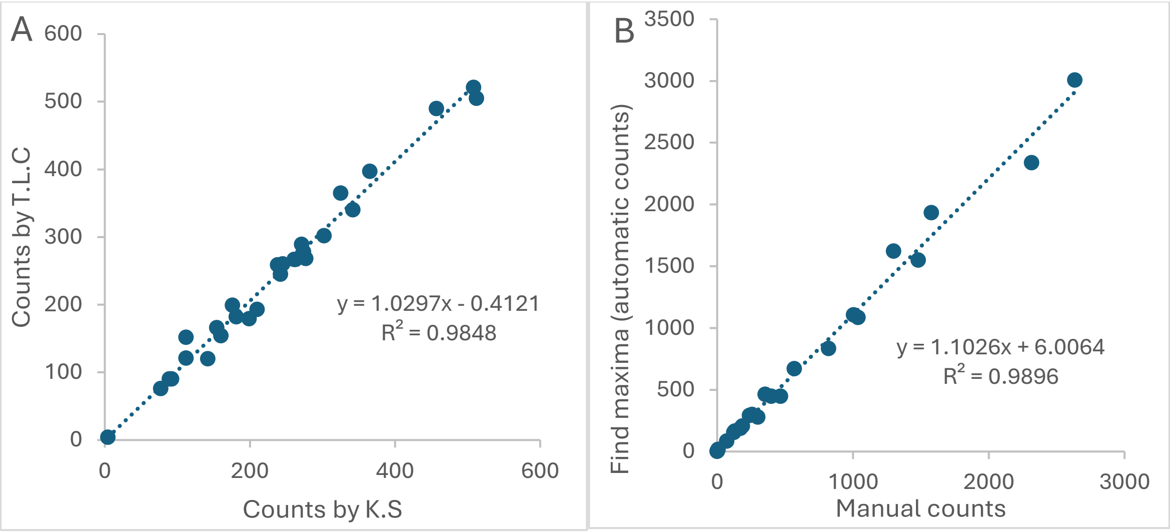


Appendix figure 5: A: high repeatability score of $R^2$ = 0.98 between two analysts, when counting sperm numbers stored in odd-numbered females after 24 hours, using the cell counter plugin. B: high repeatability score of $R^2$ = 0.98, between sperm numbers when sperm are manually counted using the cell counter plugin, versus estimated using the find maxima plugin, for sperm transferred to even-numbered females.

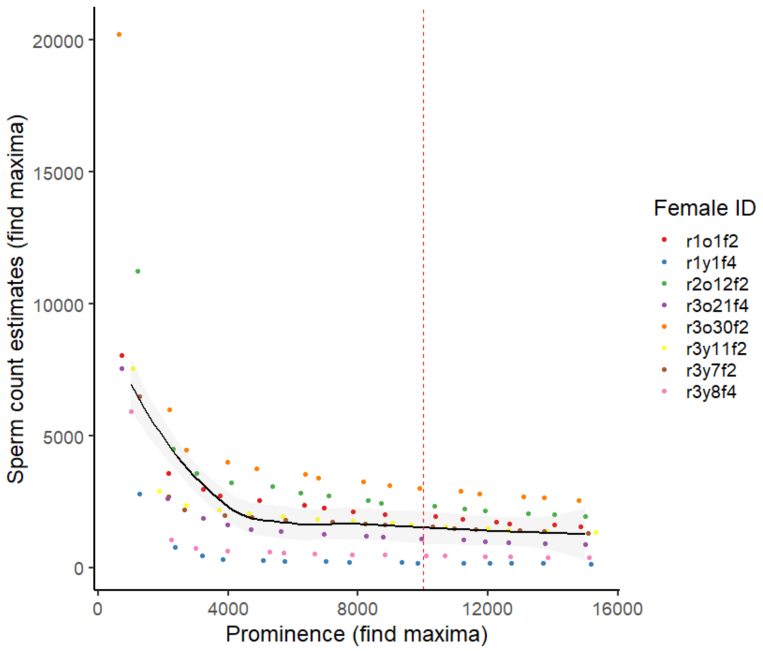


Appendix figure 6: prominence on find maxima chosen as 10000, when estimating sperm stored in females, because this is where the power function relationship between prominence and estimates of sperm number by the find maxima plugin, became linear and flat based on visual inspection.

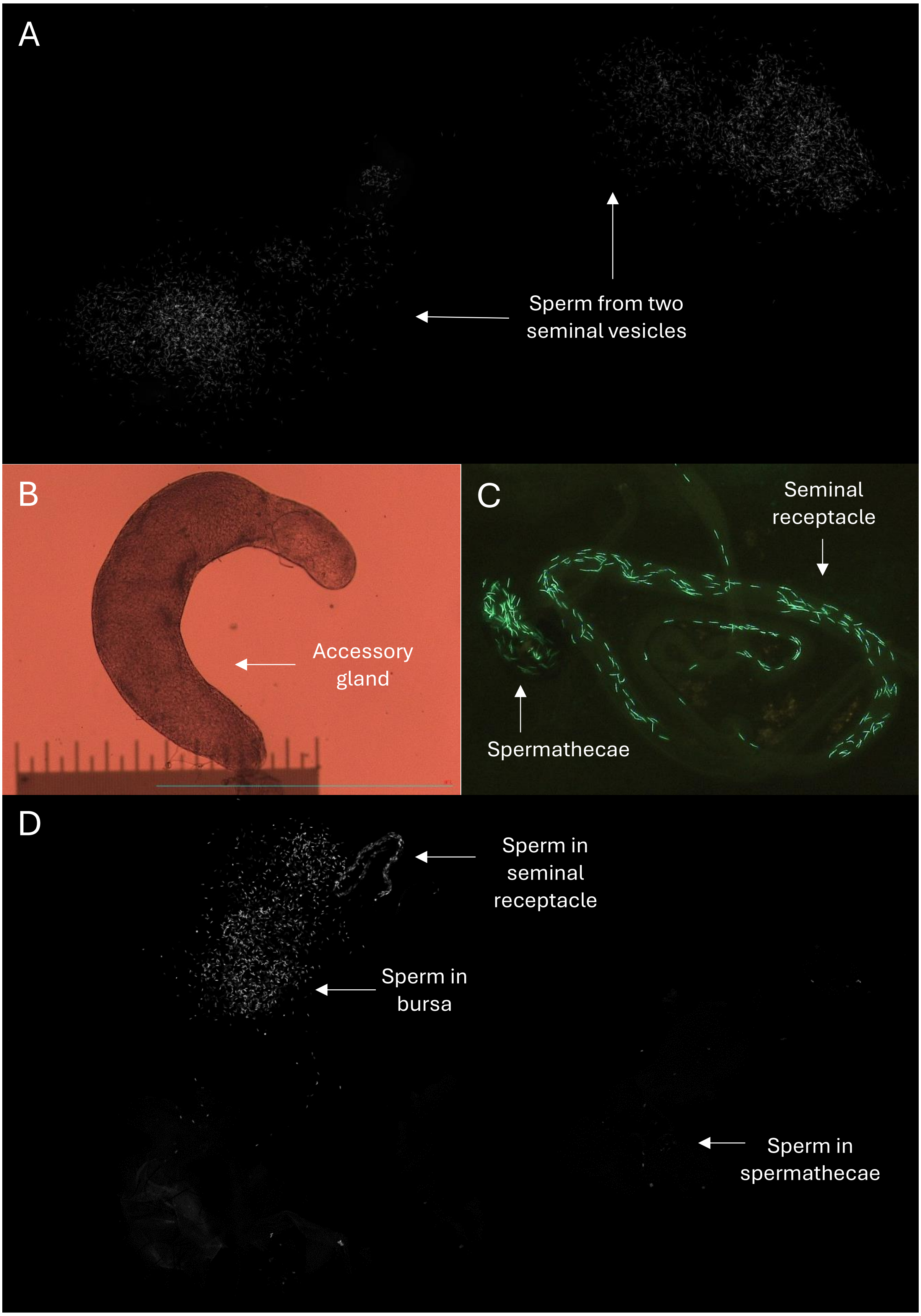


Appendix figure 7: Examples of images from dissected individuals in Experiment B. **A.** sperm in seminal vesicles of males. **B.** accessory gland in male. **C.** sperm in long-term storage organs of even-numbered female after 24 hours. **D.** sperm transferred to odd-numbered female.
